# Human primed and naïve PSCs are both competent in differentiating into bona fide trophoblast stem cells

**DOI:** 10.1101/2022.05.20.492766

**Authors:** Sergey Viukov, Tom Shani, Jonathan Bayerl, Daoud Sheban, Yonatan Stelzer, Noa Novershtern, Jacob Hanna

**Affiliations:** The Department of Molecular Genetics, Weizmann Institute of Science, Rehovot 7610001, Israel; The Department of Molecular Cell Biology, Weizmann Institute of Science, Rehovot 7610001, Israel

## Abstract

Cells of the trophoblast lineage constitute the major part of placental tissues in higher mammals. Recent derivation of human trophoblast stem cells (TSC) from placental cytotrophoblasts (CT) and from human naïve PSCs opens new opportunities for studying development and function of human placenta. Several recent reports have suggested that naïve human PSCs retain an exclusive potential to give rise to bona fide TSCs. Here we report that inhibition of TGFβ pathway and avoiding WNT stimulation, leads to direct and robust conversion of primed human pluripotent stem cells into TSCs. Systematic side by side comparative analysis showed that the latter are equivalent to previously derived TSC lines. Primed PSC derived TSC lines exhibit self-renewal, are able to differentiate into the main trophoblast lineages, and present RNA and epigenetic profiles that are indistinguishable from the TSC lines derived from placenta or naïve PSCs. Our findings underscore a residual plasticity in primed human PSCs that allows converting directly into pre-implantation extra-embryonic cell lineages.

**Highlights:**

- Primed human PSCs readily convert into TSCs upon inhibition of TGF pathway
- CHIR inhibits conversion to TSC in primed but not in naive hPSCs
- Primed human PSC derived TSCs line are indistinguishable from placental and naïve derived TSCs
- YAP is sufficient for TSC induction from hPSCs and necessary for TSC maintenance.

## Introduction

Mammalian placenta is an organ that mediates interaction between mother and fetus. It supplies the fetus with nutrients and oxygen, secretes hormones, controls maternal tissues remodeling and thus ensures the normal pregnancy progress (Turco and Moffett, 2019). Cells of the trophoblast lineage, constituting significant part of the placenta, originate from trophectoderm – an outer layer of preimplantation embryo (James et al., 2012). Segregation of trophectoderm from the Inner Cell mass (ICM) establishes the earliest cell fate decision in the development time course (Niakan et al., 2012).

At embryo implantation stage the trophoblast cells penetrate endometrium, and subsequently form the scaffold of placental villi **(Figure 1a**) (Latos and Hemberger, 2016). The inner part of this scaffold consists of undifferentiated, mononuclear cytotrophoblast cells (CT), which proliferate and can differentiate into two main types of trophoblast lineage: First, fused CT cells form the outermost layer of the villi - multinuclear syncytium trophoblast (STB) that ensures nutrient exchange between the fetus and the mother. These cells also secrete hormones like Chorionic Gonadotropin (hCG) and progesterone. Second, Extravillous Cytotrophoblast (EVT) that originate from the CT cells positioned at the distal part of the villi. These cells penetrate endometrium and maternal blood vessels remodeling the latter (James *et al*., 2012). One of the key regulators of the trophoblast lineage induction is YAP (Yes-associated protein). It is believed to be the key trigger for the trophectoderm specification of the outer cells of early mouse blastocyst (Rossant and Tam, 2009). Later in development it supports proliferation and expression of stemness genes in CT cells and prevents their differentiation (Meinhardt et al., 2020).

**Figure 1.**
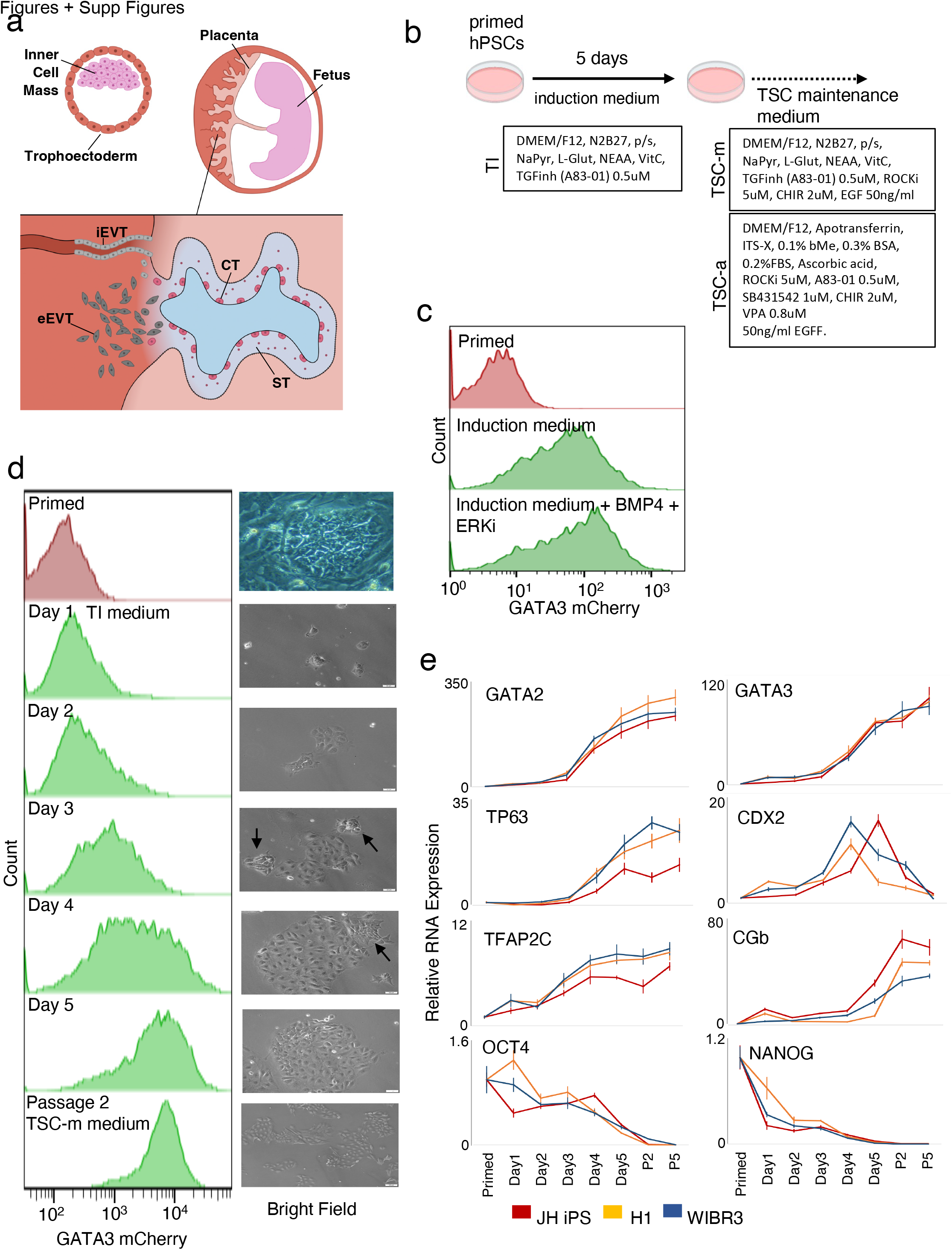
TSC markers are upregulated following TSC induction of primed hPSCs. (a) The placenta is developed from Trophoectoderm (TE). The inner part of the placenta villi consists of Cytotrophoblast (CT), which can fuse and generate Syncytiotrophoblast (ST), or exit the villi to generate Extravillous Cytotrophoblast (EVT) that penetrate endometrium (eEVT) and maternal blood vessels (iEVT). (b) Experimental flow: human pluripotent stem cells (ESCs and iPSCs) were maintained in TI medium for 5 days, and then transferred into TSC-a or TSC-m maintenance medium. (c) Levels of GATA3-mCherry reporter, as measured in WIBR3 hPSC line, maintained in primed conditions (top panel), three days after seeding in TI medium (middle panel) or in TI medium+BMP4+ERK inhibitor. (d) Left: Levels of GATA3-mCherry reporter, as measured in WIBR3 hESC line, maintained in primed conditions (top panel), during maintenance in TI medium for 5 days, and after passage in TSC-m medium. Right: Bright field image of the cell culture in the indicated time points. Arrows indicate disappearing hES-like cells. (e) Expression of selected markers in cells maintained in TI medium for 5 days, and in TSC-m medium for 2 and 5 passages. Cells lines are human iPSCs and human ESCs (H1 & WIBR3). (n=4)

Derivation of Trophoblast Stem Cells (TSC) from the 1st trimester placenta gives opportunity to study aspects of placental development and function in vitro (Okae *et al*., 2018). Particularly, TSC derived from human induced pluripotent stem cells (iPSC) from different genetic backgrounds can help model placental developmental complications.

Naïve embryonic stem cells model the ICM cells of preimplantation blastocyst, while primed embryonic stem cells model early postimplantation epiblast (Weinberger et al., 2016). These two cell states can be maintained indefinitely in vitro, and switch by changing of growth conditions, affecting several aspects such as X chromosome reactivation and the ability to contribute to chimera (Bayerl et al., 2021; Guo et al., 2017; Theunissen et al., 2014). Conversion of human pluripotent stem cells (hPSC, including both embryonic stem cells and induced stem cells) into TSC was first achieved in vitro from naïve hPSCs by applying TSC maintenance medium (Cinkornpumin et al., 2020; Dong et al., 2020; Liu et al., 2020; Lo et al., 2021; Guo et al., 2021), but not from primed hPSC state. This is in line with the ability to convert mouse naïve, but not primed, embryonic stem cells to mouse TSC (Blij et al., 2015), and is regarded as reflection of developmental proximity between mammalian ICM (modeled by naïve ESCs) and trophoblast lineage. These studies have promoted the notion that TSC differentiation potential is retained exclusively in human naïve, but not primed PSCs. However, a recent report (Wei *et al*., 2021) describes direct conversion of human primed PSCs to TSC by BMP4 stimulation. However, Polo group (Liu et al., 2020) described the sporadic isolation of putative TSC-like cells from primed PSCs, however those were not comparable to embryo derived- and naïve PSC derived TSCs based on RNA-seq clustering analysis. Thus, it remains to be proven whether the latter success (Wei *et al*., 2021) in deriving TSCs from primed PSCs was sporadic and no work has compared side-by-side TSC quality and characteristics when derived from isogenic naïve and primed PSCs. Moreover, the difference in TSC competence and signaling from these two cell states has not been resolved so far.

Here we report a novel protocol for direct and robust conversion of primed hPSCs to cell lines sharing properties, transcription and chromatin profiles with human placenta derived TSC and human naïve derived TSCs. We found that inhibition of TFGb pathway is necessary and sufficient to achieve this conversion. We have also found that addition of GSK3β inhibitor CHIR99021 (CHIR) inhibits the conversion. In addition, we show that activation of YAP, the major factor of Hippo signaling pathway can replace TGF inhibition in the conversion protocol, while YAP knockout hPSCs can show some TSC markers upregulation, but fail to proliferate, thus making YAP sufficient for this conversion, and indispensable for human TSC maintenance from both primed and naïve conditions.

## Results

### Derivation of iTSC lines from primed hPSCs

In order to study conversion of primed/conventional and naïve hESC and hiPSC lines into trophoblast lineage, a reporter for GATA3, which is highly expressed both in trophectoderm and in the first trimester placenta (Petropoulos et al., 2016; Soncin et al., 2018), was generated using CRISPR-Cas9 strategy **(Supp Figure 1)**. Shortly, P2A-mCherry reporter was inserted at the 3’ end of GATA3 coding sequence in WIBR3 hESC line, which was maintained at primed conditions **(Table 1)** from the moment of derivation (Lengner et al., 2010). We then attempted to induce these human primed GATA3-reporter cells (W3GC) to become TSCs, using TSC Induction medium (TI), which includes N2B27 & TGF inhibitor (A83-01, **Table 1)**. Three days after seeding those cells in TI medium **(Figure 1b)**, GATA3 reporter was robustly upregulated **(Figure 1c)**, indicating trophoblast program induction.

**Table 1:**
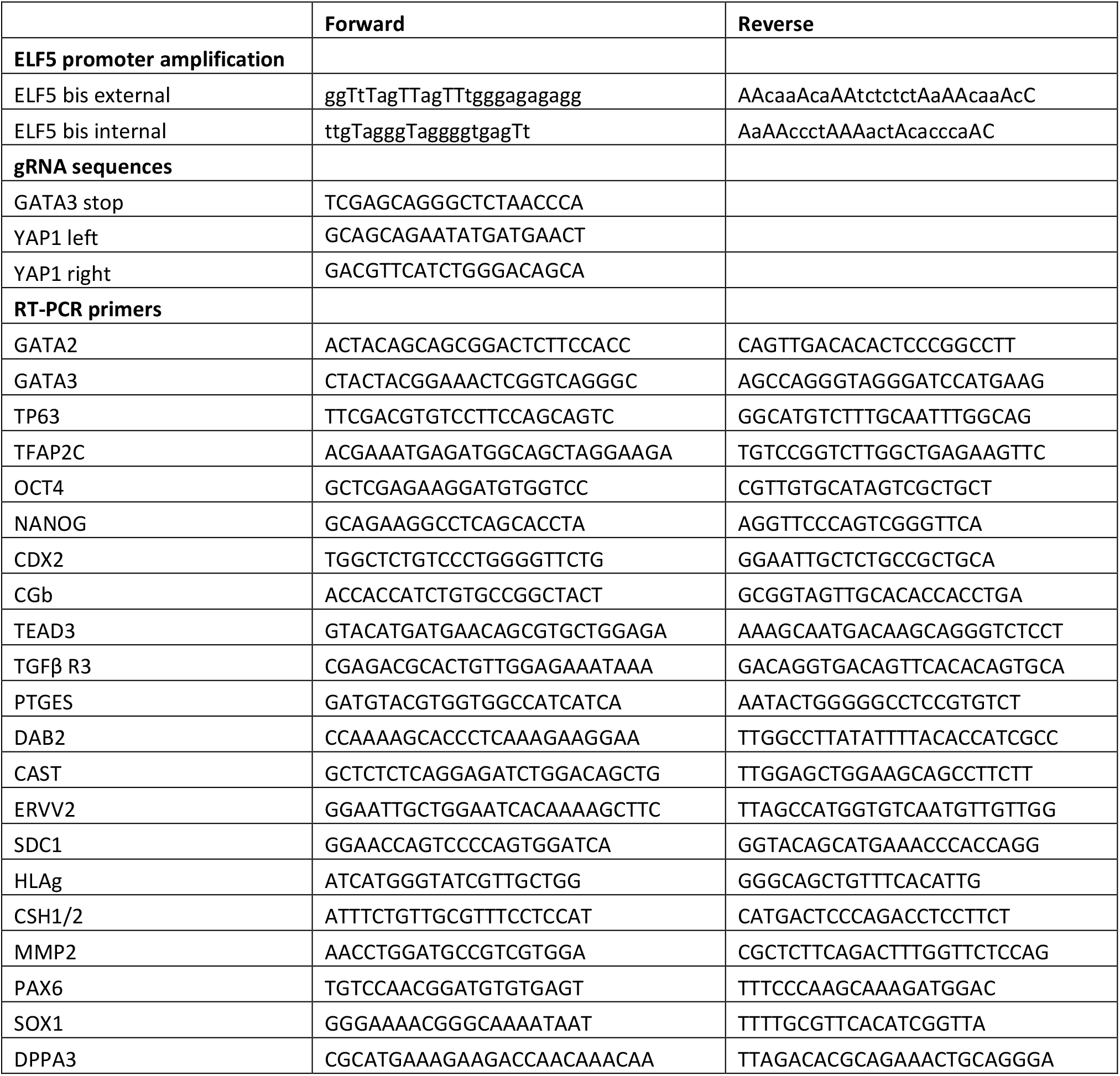
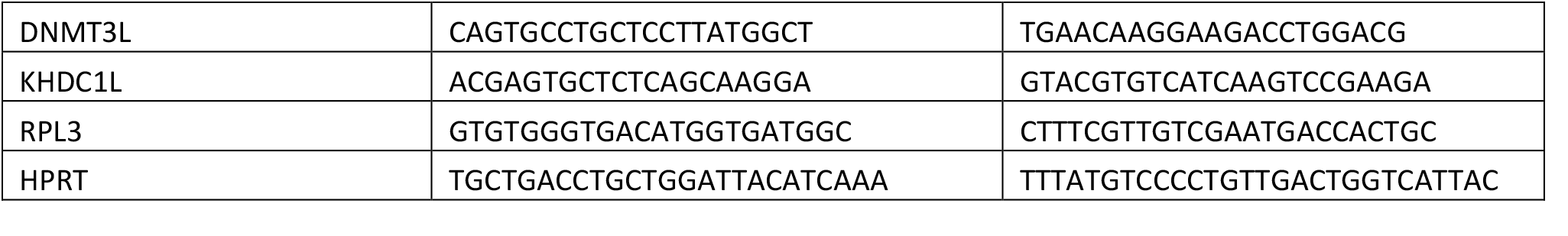
primer list

Next, W3GC, H1 hESC and JHiPS primed cells were seeded into TI medium and expression of trophoblast markers was monitored daily. mCherry upregulation was apparent 24 hours after induction, while by day 5 more than 90% of the cells expressed it at high level **(Figure 1d)**. Additional trophoblast markers GATA2, TP63, TFAP2C, CDX2 and CGb gradually increased and pluripotent markers OCT4 and NANOG decreased during five days of induction **(Figure 1e)**, indicating that the cells were effectively converted to the trophoblast lineage. The morphology of the cells also changed gradually. For example, by day 4 **(Fig 1d, right panel**) most of the cells have already acquired cobble-stone flatten TSC-like appearance, while some small groups of cells still looked more like primed hESCs. By day 5 the latter cells were not visible. Despite not using ROCK inhibitor in this experiment, we have not noticed significant cell death in the cultures, and the colonies rapidly increased in size.

After 5 days in TI medium, the cells grew confluent and were passaged to Collagen-IV-coated plates into TSC maintenance medium (TSC-m, including N2B27, A83-01, EGF, GSK3beta inhibitor CHIR99021 (CHIR) and ROCK inhibitor Y27632) **(Figure 1b)**. After one or two passages in TSC-m medium, all cells showed homogeneous TSC morphology and GATA3 expression **(Figure 1d)**, while TSC markers persisted their induction **(Figure 1e)**. One interesting exception was CDX2 which peaked around day 5 of induction of primed hPSCs and then was repressed by passage 1-2. This is in line with the previous reports (Dong *et al*., 2020), according to which, CDX2 is similarly transiently upregulated during naïve hPSC to TSC conversion and cannot be maintained in any of previously described human TSC lines.

TSC state remained stable after prolonged passaging. High expression of trophoblast markers was detected by RT-PCR after 10-15 passages in TSC-m maintenance medium **(Figure 2a)**. Cells were stained positive for TFAP2C, GATA3 and KRT7 (CK7 protein) after 15 passages **(Figure 2b)**, and trophoblast marker ITGA6 was also highly expressed as detected by FACS **(Figure 2c)**. Cells could be kept for at least 30 passages without changes in their morphology and markers, showing no changes in cell identity **(Figure S2a, b)**. Using the same protocol, we have converted to TSCs other published primed/conventional hPSC lines, which have never been maintained in other than primed conditions (H9 ESC, WIBR1 ESC, WIBR2 ESC, JH22 iPSC). These TSC lines have shown the same morphology and markers expression as the ones converted from WIBR3 background and as TSC lines derived from human placenta and from blastocyst **(Figure 2a, S2c)**, indicating that the identity of *in vitro* obtained TSCs is independent in the genetic background of the cells. These results exclude the possibility that the ability to obtain TSCs from primed hPSCs is a sporadic and/or cell line specific phenomenon. We labeled the cells as primed state derived TSCs – pdTSC.

**Figure 2.**
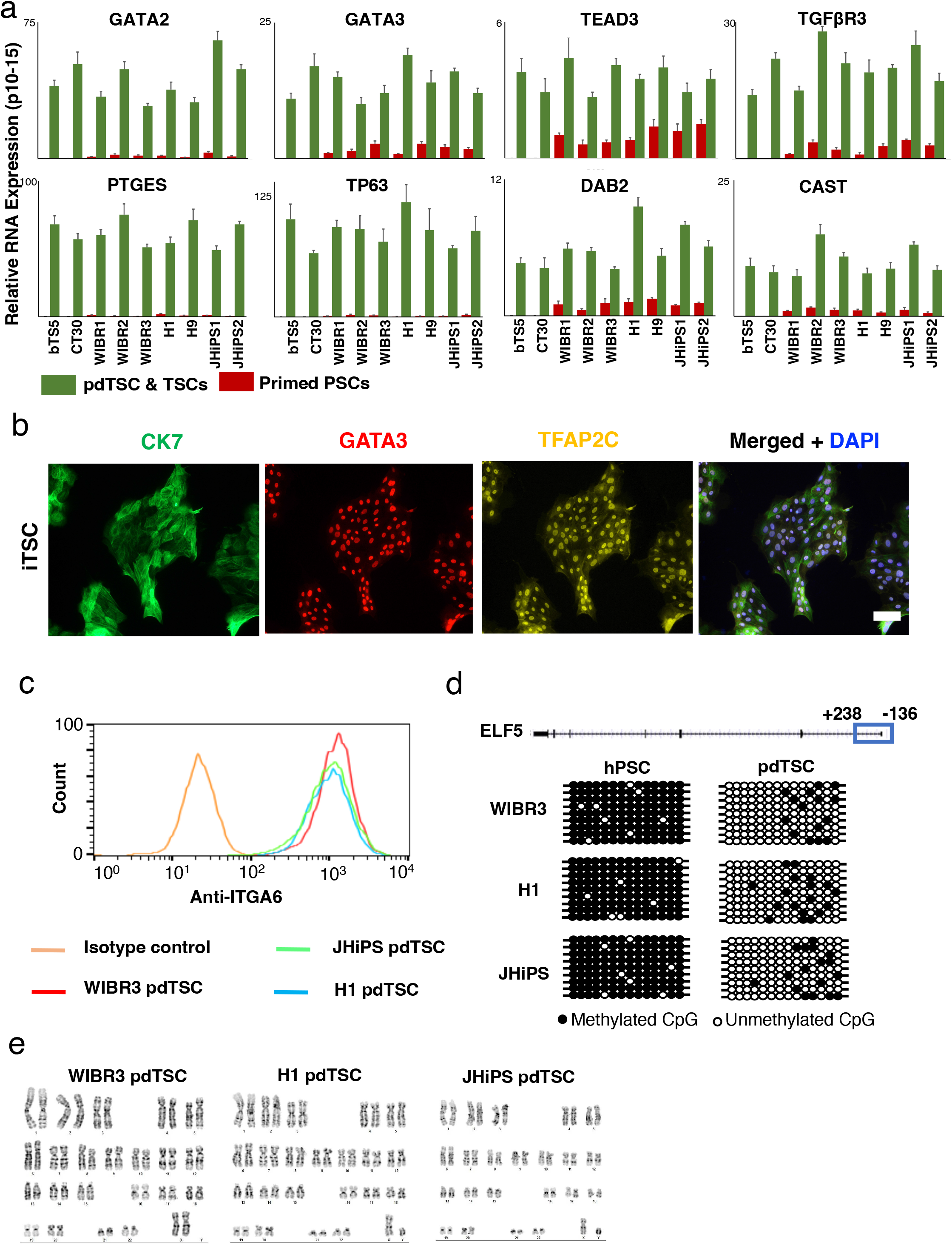
Cells retain TSC identity after prolonged time. (a) Expression of trophoblast markers as measured by RT-PCR in pdTSCs that were maintained for prolonged time (10-15 Passages) in TSC-m medium, or in primed hPSCs, compared to native TSC lines bTS5 and CT30. (n=3) (b) Immunostaining of trophoblast markers CK7, GATA3 and TFAP2C in cells that were maintained for prolonged time (15 Passages) in TSC-m medium. (c) FACS analysis of H1, WIBR3 and JHiPS pdTSCs, measuring anti-ITGA6 or IgG (negative control), after 15 passages in TSC-m medium. (d) Methylation levels of ELF5 promoter, as measured in primed hPSC (left) and in pdTSC (right). (e) Karyotype analysis for H1, JHiPS and WIBR3 pdTSCs.

We next moved to conduct in depth validation of pdTSCs. Demethylation of ELF5 promoter is considered to be an important marker of human TSCs (Hemberger et al., 2010). Methylation status of ELF5 promoter was estimated in three pdTSC lines and in their parental primed hPSC: While ELF5 promoter was highly methylated in hPSCs (>94% methylation) the methylation was lost in all iTSCs measured **(Figure 2d)**. Importantly, the karyotype of pdTSC remained stable even after 20-25 passages, with no apparent chromosomal aberrations **(Figure 2e)**.

We note that the TSC-m medium described herein is similar to conditions that were described earlier (Okae *et al*., 2018) (TSC-a, **Table 1, Figure 1b**), but lacks HDAC inhibitor (VPA) and is based on B27 supplement, while the conventional growth medium contains low concentration of FBS. We could similarly derived pdTSCs from the same primed hPSCs by using TSC-a medium described previously by Okae et al. Comparing the pdTSCs expanded in TSC-m and TSC-a media conditions, similar marker expression was detected **(Figure S2c)**, however TSC-a cells had lower growth rate **(Figure S2d)** and higher expression of EVT and STB differentiation markers (HLAg and hCG, **Figure S2e**), therefore for routine maintenance, TSC-m conditions were used.

### pdTSC lines share transcriptional profile and chromatin configuration with other TSC lines

Consistent with previously described papers (Cinkornpumin et al., 2020; Dong et al., 2020; Io et al., 2021; Guo et al., 2021), isogenic naïve hPSC expanded in HENSM conditions (Bayerl et al., 2021), generated TSC lines, termed ndTSC-naïve state-derived TSCs, from multiple cell lines (H1, WIBR3 and JH22-iPSC).

Next, RNA-seq profiles of H1 and WIBR3 derived pdTSC and ndTSC were compared to previously described embryo-derived TSC (edTSC) lines derived from human blastocyst and first trimester placenta (Okae *et al*., 2018). All lines were maintained in both TSC-m and TSC-a conditions. In addition, profiles were compared to primed hPSCs and human fibroblasts, and to previously published ndTSC and hPSC samples (Dong *et al*., 2020; Liu et al., 2020). All TSC lines clustered together and separately from hPSCs and fibroblast **(Figure 3a)**, showing the transcriptional difference between derived TSC and other cell types. Overall, 894 genes were upregulated in TSC compared to hPSC, and were highly enriched for placental genes, including well known TSC markers such as GCM1, CGA, CGB5, ERVFRD-1, KRT7, GATA2, GATA3 & ELF5 **(Figure 3b, Figure S3)**. Cannonical pluripotency genes such as OCT4, SOX2, NANOG & DPPA4 were downregulated in all TSCs as expected **(Figure 3b)**.

**Figure 3.**
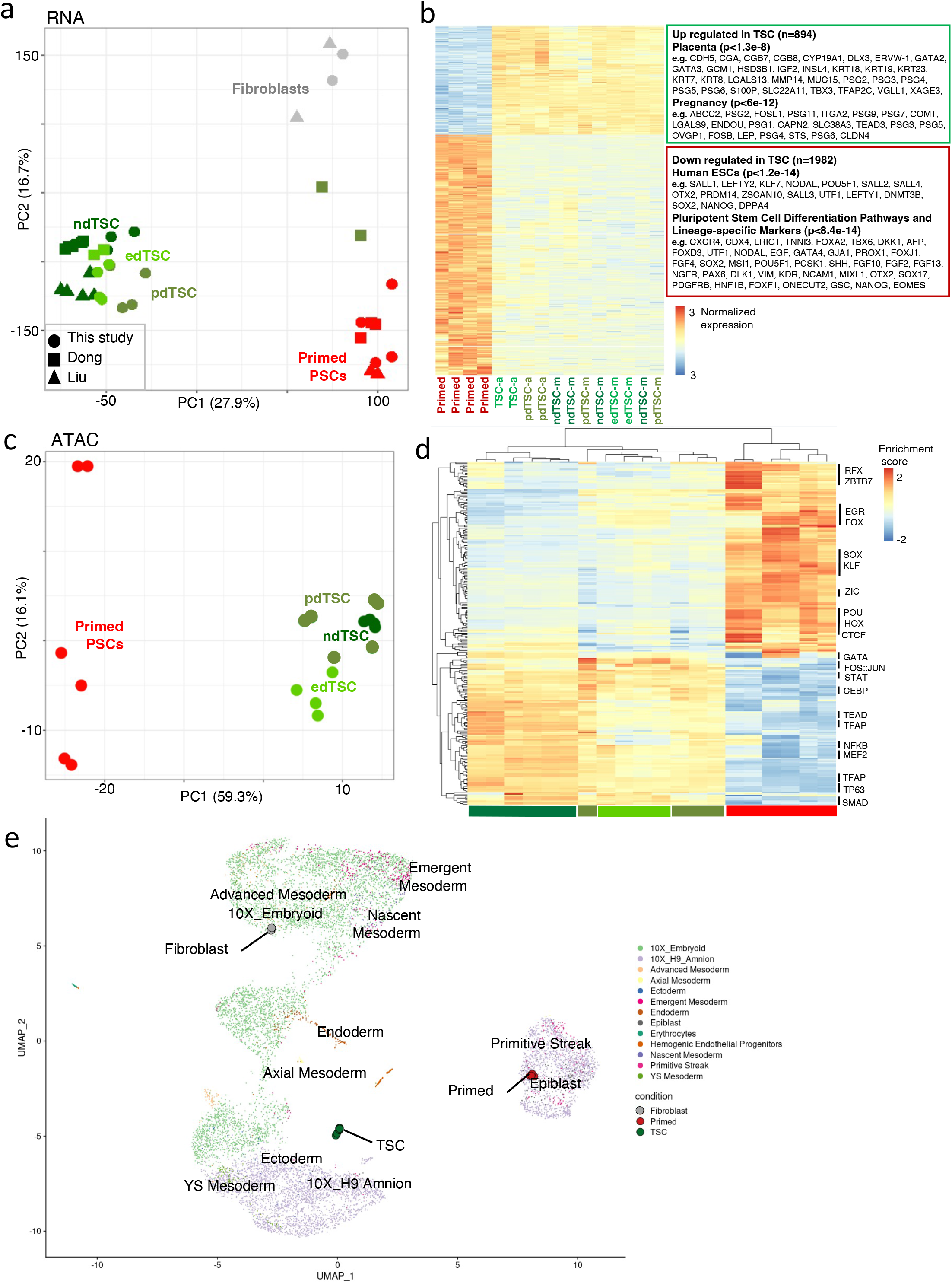
pdTSC transcriptional profile is indistinguishable from naïve and embryo derived TSCs. (a) Principal component analysis of RNA-seq profiles from pdTSCs, ndTSCs, primed hPSCs and Fibroblasts alongside previously published datasets (Dong et al., 2020; Liu et al., 2020), showing that pdTSCs cluster with edTSC, and both are closer to EVT and ST than to hPSCs or fibroblasts. pdTSC & ndTSC were derived from H1 and WIBR3 primed and naïve cells. For Naïve lines the derivation was performed by direct application of TSC-m medium and through induction step. (b) Differentially expressed genes between edTSC/pdTSC/ndTSC and primed hPSCs, 894 genes were upregulated in TSC, enriched for placenta related gene signatures. 1981 genes were downregulated in TSC, enriched for embryonic stem cells gene signatures. Selected genes are highlighted. The complete DEG lists and enrichment are available in **Supp Table 3**. (c) Principal component analysis of pdTSCs, ndTSC, TSCs and naïve and primed hPSCs samples, which is based on motifs enriched in accessible chromatin regions (ChromVar), showing that pdTSCs cluster with TSC, away from primed hESCs. (d) Motif enrichment in ndTSC, pdTSC, edTSC, and primed hPSC samples, calculated from ATAC-seq data using ChromVar. Selected motif families are highlighted. (e) Projection of RNA-seq data onto published datasets (Tyser et al., 2020; Zheng et al., 2019), showing that while primed hPSCs and fibroblast fall within the expected cell types (Epiblast and Advanced mesoderm, respectively), pdTSC/ndTSC/edTSC are not projected on amnion cell type.

Next, Chromatin configuration was measured in TSC and hPSC samples using ATAC-seq. Motif enrichment analysis of open chromatin regions, clustered the samples into distinct clusters **(Figure 3c)**, showing that the difference between TSC and other cell types is robust and rewired in the chromatin configuration. The top motifs that are associated with all TSC samples include known TSC regulator families such as GATA, TFAP, TEAD, CEBP and FOS-JUN, while the top motifs that are associated with hPSC samples include OCT4 (POU), SOX and KLF **(Figure 3d)**. Both observations are well coordinated with the expression and function of these transcription factors **(Figure S3a)**.

A recent paper (Zhao et al., 2021) has described principles and markers to be used to avoid confusion between identifying placental vs. amniotic human cells. To rule out the possibility that our pdTSC cells are amniotic, their RNA-seq profiles were projected on reference single-cell RNA-seq datasets (Tyser et al., 2020; Zheng et al., 2019). While fibroblasts and primed hESCs were projected on their corresponding in-vivo counterpart **(Figure 3e)**, pdTSCs were not projected on amnion cells. In addition, amnion markers such as BAMBI, ISL1 and IGFBP3 were not upregulated in pdTSC compared to edTSC or to primed hPSCs **(Figure S3b)**, thus ruling out the possibility that the cells are amniotic. Overall, these results show that pdTSC share transcriptional and chromatin profiles with previously published ndTSC and with edTSC derived directly from the placenta, therefore authentically represent TSC lines.

### pdTSCs can differentiate into the main trophoblast lineages and form lesions upon injection into NOD-SCID mice

A key property of TSCs is their ability to differentiate to the main trophoblast lineages (STB and EVT), to which they give rise in vivo. To test whether our pdTSCs can generated STB, as previously described for edTSC and ndTSCs (Cinkornpumin *et al*., 2020; Dong *et al*., 2020, Okae *et al*., 2018), pdTSCs were treated with STB differentiation protocol (Okae *et al*., 2018), and after 6 days formed multinucleated synthitium, with a typical multinucleated structure **(Figure 4a)**. Strong STB markers upregulation was detected by RT-PCR and immunostaining (CGb, ERVV2, SDC2, **Figure 4b, c)**, and high levels of hCG hormone, which is typically secreted by STB, was identified by ELISA in the medium **(Figure 4d)**. To prove that these multinucleated structures are formed through cell fusion, pdTSC were marked by constitutive GFP or tdTomato **(Figure 4e)**. GFP and tdTomato expressing pdTSC were mixed and cultured in STB conditions. While cells that remained in TSC medium formed separate clusters of red and green cells, the multinucleated ST cells expressed both markers thus proving cells underwent fusion **(Figure 4e)**.

**Figure 4.**
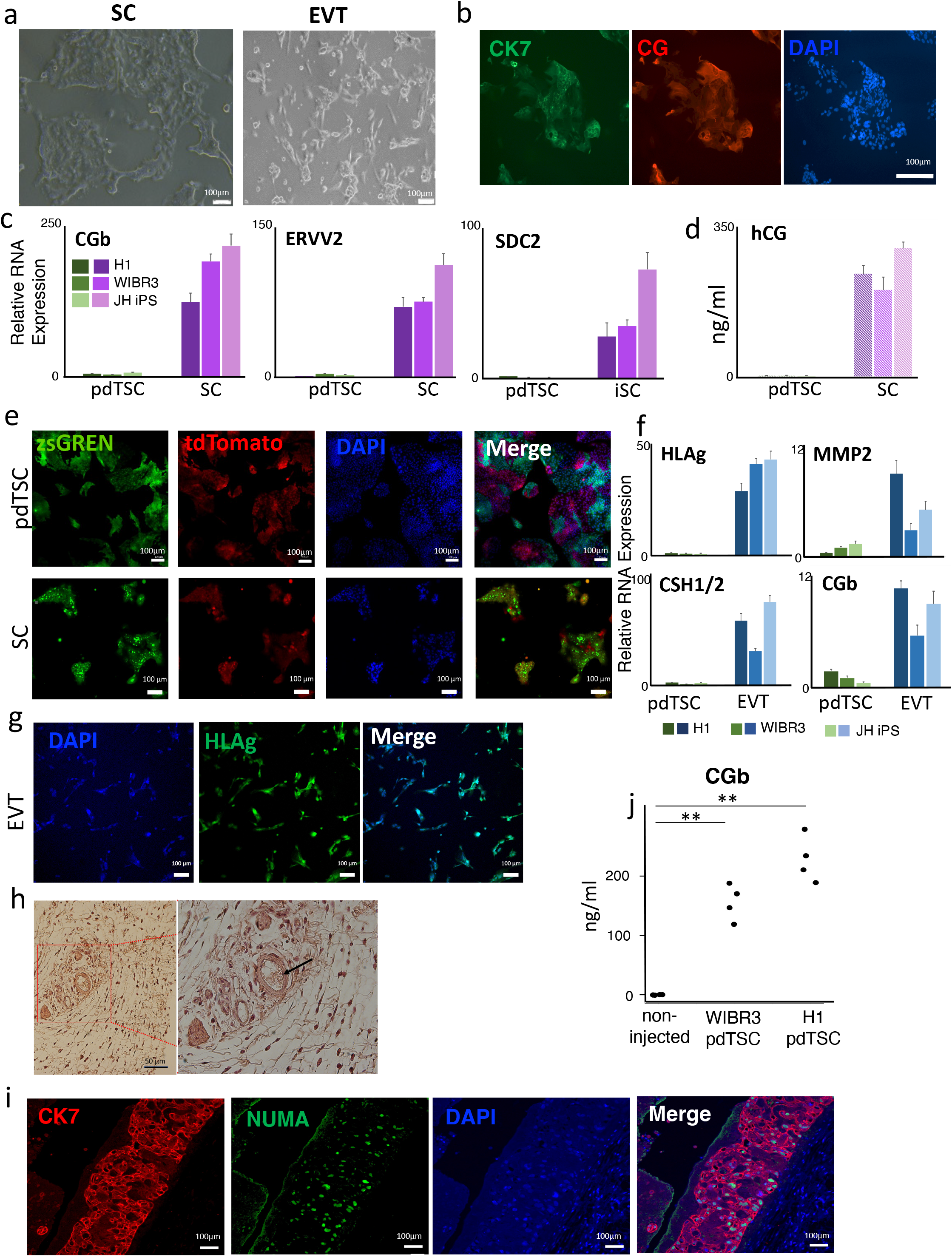
Formation of placental lineages iST and iEVT from pdTSC. (a) Representative images of induced Syncytium Trophoblast (left) and induced Extravillous Cytotrophoblast (right), generated from pdTSC at the end of the 6 days differentiation protocols. (b) Immunostaining of ST markers CK7 and CG, in ST cells derived from pdTSC. (c) Relative expression of ST markers CGb, ERV-V2 and SDC2, measured by RT-PCR. (n=3) (d) Levels of hCG secreted hormone, measured by ELISA. (n=3) (e) A mixture of GFP and tdTomato expressing pdTSCs after treatment with ST differentiation medium (bottom), and without treatment (top). (d) Relative expression of EVT markers HLAG, MMP2, CSH1/2 & CGb measured by RT-PCR. (n=5) (e) Immunostaining of EVT marker HLA-G in EVT cells derived from pdTSC. (f) H&E staining of the pdTSC-lesions formed in male NOD-SCID mouse. Arrow indicates blood filled lacunae, surrounded by trophoblast cells. (g) Immunostaining for CK7 and NUMA in iTSCs-formed lesions. (h) hCG levels in serum of host male mouse that was injected with pdTSC derived from WIBR3 and H1 hES or non-injected control. ** t-test p-value <0.0012

To show differentiation of pdTSCs into EVT lineage, cells were differentiated using conventional protocol (Okae *et al*., 2018). The resulting cells expressed high levels of EVT markers such as HLA-G, MMP2, CSH1/2 & CGB **(Figure 4f)** and stained positive for HLA-G protein **(Figure 4g)**. Lastly, pdTSCs were able to form lesions following injection into mice: H1 and WIBR3 derived iTSCs were injected to male NOD-SCID mice. After 7 days, 3-7mm lesions were observed **(Figure 4h)**. This lesions had necrotic middle region, surrounded by CK7 positive CT-like cells (**Figure 4h,i)**. Other cells morphologically resembled STB cells and contained blood-filled lacunae **(Figure 4h)**. To prove that the injected cells are the source of the lesions, the samples were co-stained for specific human antigen NUMA **(Figure 4i)**. Host mouse serum contained significant amount of hCG **(Figure 4j)** in correspondence with the previous reports (Okae *et al*., 2018; Turco et al., 2018), indicating the injected cells retain trophoblast properties.

### Huma naïve PSCs are not more susceptible to generate iTSCs than primed PSCs

Recent reports claim that only naïve human pluripotent cells retain potential to convert into TSC-like cells (Cinkornpumin et al., 2020; Dong et al., 2020; Io et al., 2021; Guo et al., 2021). Since our results show that primed hPSCs can readily generate TSCs **(Figures 1, 2)**, we checked whether the cells that generate TSCs are passing transiently through a naïve state. The conversion protocol was therefore applied to WIBR3, H1 and JHiPS primed hPSC lines, and the expression of naïve markers was measured daily by RT-PCR. No significant upregulation of stringent naïve markers (DPPA3, DNMT3L, KHDC1L) was detected at any day of the induction process **(Figure 5a)**, indicating that the cells do not pass through some transient quasi-naïve state during their conversion.

**Figure 5.**
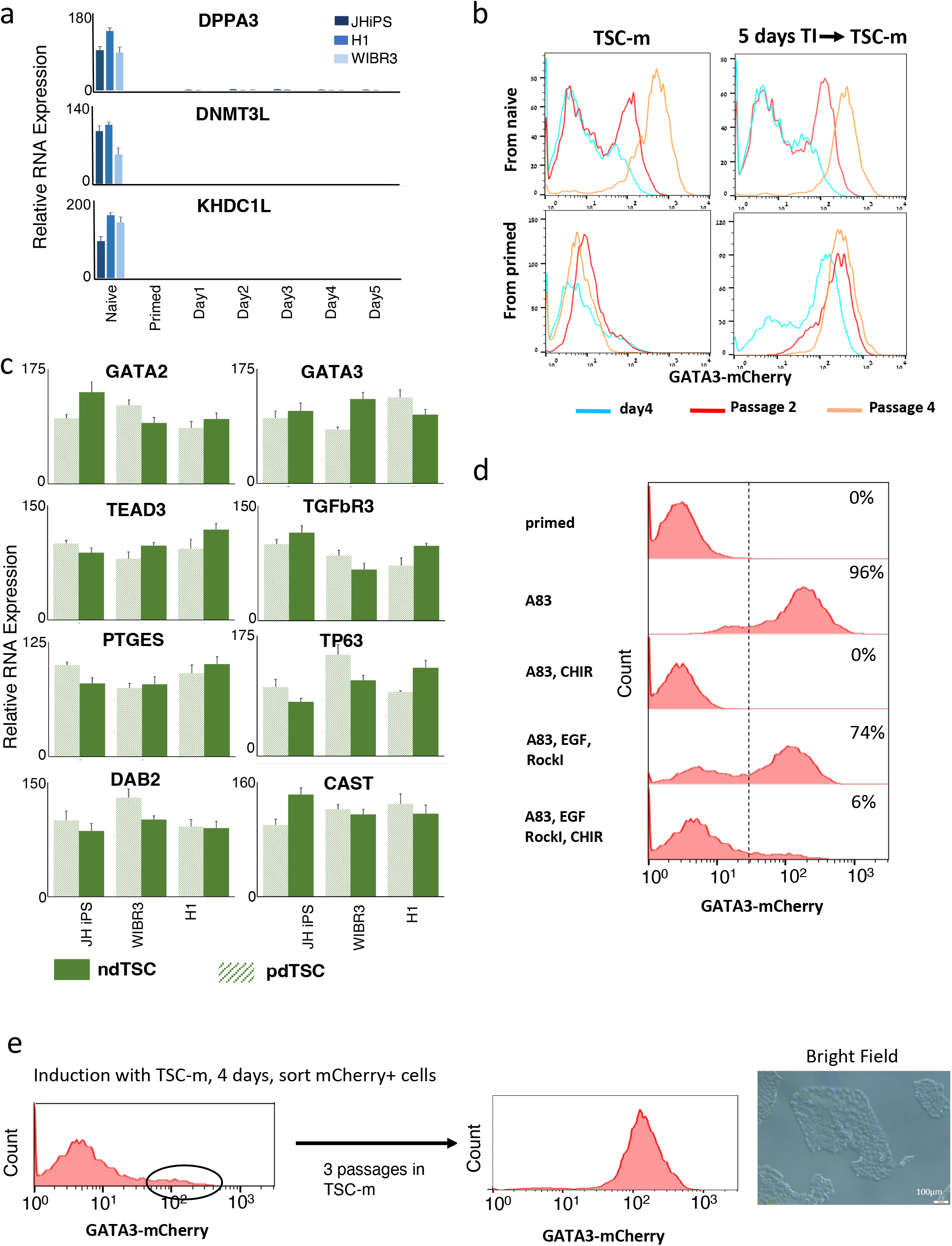
Derivation of TSCs from Naïve and Primed hPSCs. (a) Relative expression of naïve hPSC markers DPPA3, DNMT3L and KHDC1L in naïve and primed hPSCs, and during 5 days in TI medium, measured by RT-PCR. (n=3) (b) Conversion kinetics of naïve and primed GATA3-Cherry hPSCs, by direct application of TSC-m medium (left) or by 5 days treatment with TI medium followed by TSC-m medium (right). (c) Relative expression of TSC markers, measured in induced TSC which were derived from naïve or primed hPSCs of the indicated backgrounds. (n=3) (d) GATA3-mCherry primed hPSC treated with N2B27 with indicated compounds added, showing abrogation of GATA3 expression by CHIR. Note a 6% GATA3-cherry fraction in the cells treated with TSC-m (lower panel). (e) Lower panel from (d). mCherry positive fraction is sorted out and gives rise to *bona fide* TSC culture.

As previously published TSC conversion protocols applied TSC maintenance media directly on naïve human ESCs, without using a prior inductions step as devised in our protocol described herein (Cinkornpumin et al., 2020; Dong et al., 2020; Io et al., 2021). Thus we tested the ability of hPSCs to convert into TSCs by applying alternative previously described protocols that do not use an induction step. To test this, TSC-m medium was applied directly on isogenic naïve and primed human GATA3-mCherry reporter cells, without the prior induction step. While naïve line induced GATA3-mCherry reporter expression in about 50% of the cells after 4 days, primed cells nearly failed to do so **(Figure 5b, left)**. These results corroborate the findings previously published (Dong et al., 2020).

Next, naïve and primed hPSCs were differentiated into TSCs using 5 days in TI medium and shifting to the TSC-m medium **(Figure 5b, right)**. For naïve cells, the conversion kinetics were similar using both protocols, forming up to 100% GATA3-Cherry positive population by passage

4. Interestingly however, the conversion of naïve cells using TI medium was slightly and consistently slower compared to primed: naïve cells established homogeneous ndTSC populations by passage 3-5 while TSCs derived from primed cells showed uniform GATA3 expression and *bona-fide* TSC morphology already by passage 2 **(Figures 1d, e, 5b)**. These results indicate that the inability or difficulty to obtain human TSCs is not an inherent property of the human primed state, but rather reflects a technical factor – suboptimal differentiation protocols that was not customized for human primed cells which need an induction step as delineated herein **(Figure 1b)**.

Naïve hPSC lines from 3 different backgrounds were converted into TSCs by direct TSC-m medium application, showing similar markers expression compared to their counterparts converted from primed lines **(Figure 5c)**. Some anecdotal and subtle differences in expression were observed between ndTSCs and pdTSCs, such as mild relative up-regulation of DPPA4 in ndTSCs **(Figure S3c)**. However, overall, the high similarity between the profiles proves that the induced TSCs show an authentic TSC-like signature, regardless of the origin of the cells they were derived from.

### Inclusion of GSK3i derails human primed PSCs away from the TSC lineage

The above results support the conclusion that one of the components in TSC-m, which is not found in TI, negatively influences the ability of primed PSCs to successfully convert into TSCs. Indeed, when TSC-m medium was applied directly on primed hPSCs and in accordance with the previous reports (Dong *et al*., 2020) and without a prior treatment with TI medium, the cells generated GATA3-cultures that showed homogeneous morphology and proliferated rapidly for at least 20 passages. However, these cells did not express trophoblast markers, but a mix of lineage commitment markers like PAX6 and SOX1 **(Figure S5)**.

We therefore attempted to induce TSC in primed cells cultures by applying TSC-m medium with separate addition of the TSC-m components on TI base media, and realized that addition of CHIR, GSKB3 inhibitor that leads to WNT activation, is a major inhibitor for obtaining GATA3+ cells, and that its omission from the TSC-m medium permits derivation of TSC lines from primed cells even in the presence of EGF and ROCK Inhibition without the need for TI step **(Figure 5d**). Moreover, we note that direct application of TSC-m medium on primed GATA3-mCherry cells did yield a small GATA3 positive fraction. After sorting out these Cherry-positive cells we have established normal TSC validated lines **(Figure 5e)**.

### YAP is necessary and sufficient for conversion of human ESCs into TSCs

YAP translocation to the nucleus in the morula outer cells is a crucial event in the induction of mammalian embryo trophectoderm (Rossant and Tam, 2009). Induced nuclear localization of YAP cofactor Tead4 is sufficient to convert mouse naïve ESCs into TSCs (Nishioka et al., 2009). We tested whether constitutively active YAP2-5SA (YAP*) (Zhao et al., 2007) gene overexpression, can induce TSCs from hPSC. Primed hPSCs were electroporated with YAP*-IRES-CFP, or with control IRES-CFP construct **(Fig 6a)** and seeded into N2B27 medium without TGF inhibitor. After 4 days the control cells did not express GATA3 reporter, while in the cultures transfected with YAP* nearly all the cells that expressed CFP also expressed mCherry **(Fig 6b)**. A minor CFP-mCherry+ fraction could also be observed, probably indicating the cells that were induced by transfected YAP*,but have lost the expression plasmid by the time of the measurement. This result indicates that upregulation of YAP signaling alone is sufficient to initiate conversion to TSC in minimal basal conditions.

**Figure 6.**
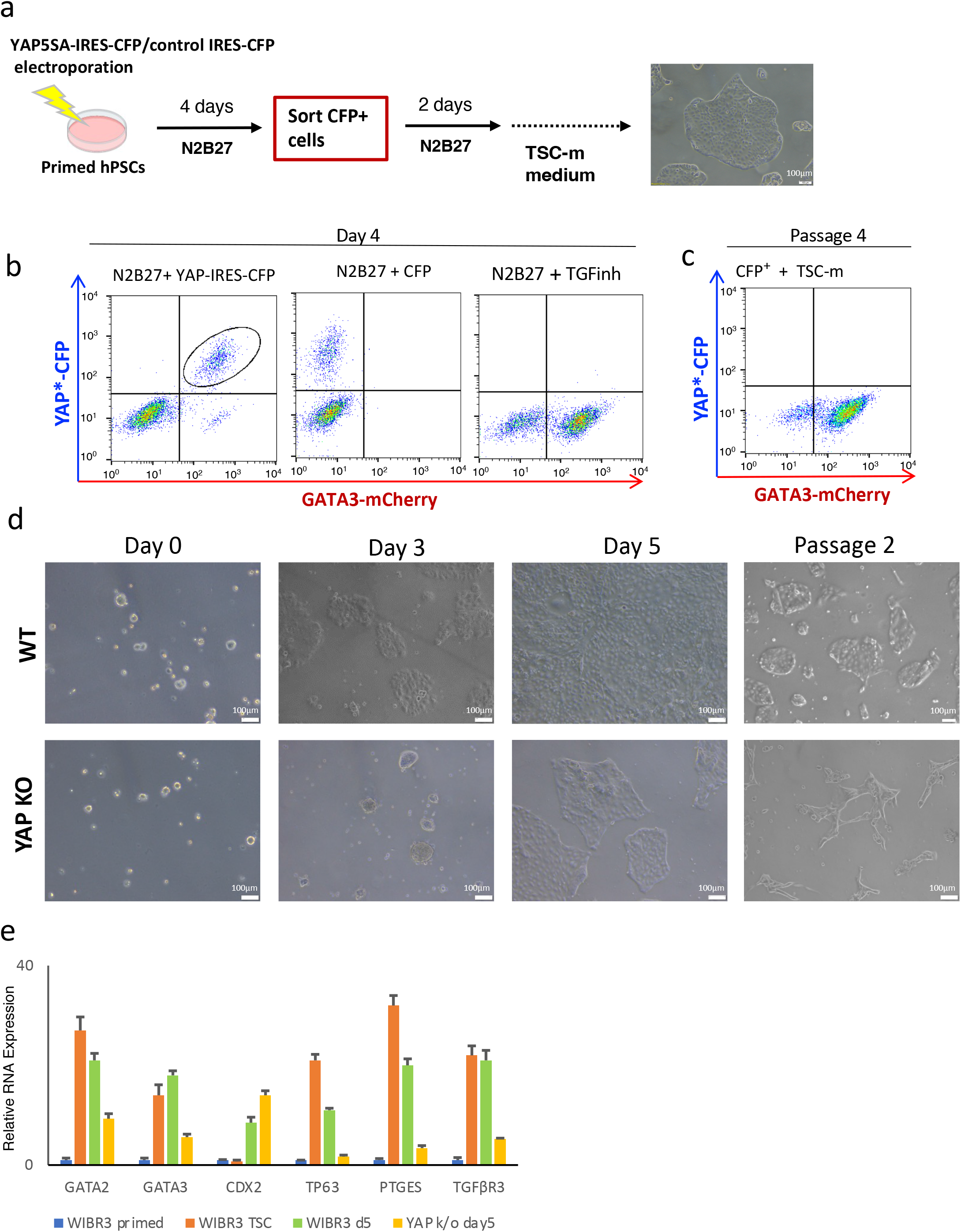
Role of YAP in the TSCs derivation and maintenance. (a) YAP* overexpression experiment scheme. (b) GATA3-mCherry reporter and CFP expression in the cells treated with N2B27 medium for 4 days. Left: Cells with YAP*-IRES-CFP construct. Middle: Cells with empty vector (negative control) Right: Cells treated with TGF inhibitor (positive control). The indicated mCherry^+^CFP^+^ fraction was sorted and maintained in TSC-m medium. (c) The sorted fraction state after 4 passages. Note that CFP is not expressed already at this time point, indicating absence of exogenous YAP*. (d) Conversion of WIBR3 WT primed hPSC (upper panel) and YAP^-/-^ hPSC. Note TSC-like morphology at day5 in YAP^-/-^ cells, however stop growing and degrade at passage2. (e) RT-PCR detection of TSC markers of WIBR3 WT and YAP^-/-^ primed hPSC after 5 days in TI medium, compared to primed hPSC and established pdTSC (both WIBR3) at passage 10. Note moderate upregulation of GATA2, GATA3, PTGES and TGFβR3 and strong upregulation of CDX2. Representative result out of 3 independent experiments (n=3).

Four days after electroporation, CFP+GATA3+ population was sorted and seeded back to N2B27 medium for two days, and then changed to TSC-m medium. The cells gave rise to self-renewing cultures with TSC morphology **(Fig 6a)**. The experiment was repeated with other primed hPSC lines (H1 and JHiPS) which do not carry GATA3 reporter, here the CFP+ cells were sorted out and gave rise to pdTSC cultures. CFP+ cells could not be detected by passage 3-4, indicates that YAP* overexpression was indeed transient **(Fig 6c)**. Importantly, all YAP induced TSC lines were readily differentiated into EVT and STB lineages, as demonstrated by markers expression **(Figure S4d-f)**, proving that that YAP signaling alone is sufficient to induce *bona fide* TSCs with adequate functional capabilities.

Next, to check whether YAP is essential for TSC generation, YAP gene was knocked-out in WIBR3 cells **(Figure S1c)**. TI medium was applied on YAP-KO cells, and while during the first 5 days they have started to show TSC-like morphology, the cell growth was impaired compared to the wild type control **(Figure 6d)**. When the cells were passaged at day 5 to TSC-m medium they stopped proliferating and eventually died. RT-PCR on the samples collected at day 5 show that some TSC markers such as GATA2, GATA3, PTGES and TGFβR3 were only mildly upregulated, TP63 was not induced **(Figure 6e)**.

## Discussion

In this study we have developed a robust protocol for conversion of primed hPSCs into TSC lines, and based on side by side comparison of isogenic lines, they were equivalent in features to TSCs differentiated from naïve hPSCs and to TSC lines derived from early placenta and preimplantation blastocyst. Those pdTSCs can differentiate into the main human trophoblast lineages – EVT and STB, and they can form KRT7 positive lesions in the NOD-SCID mice, while rising the concentration of HCG hormone in the injected male mice blood. Furthermore, by performing ATAC-seq and RNA-seq we have demonstrated transcriptional and epigenetic similarity between pdTSC and all previously generated TSC lines.

Previous reports described derivation of TSC lines from naïve hPSC by simply switching the growth medium from ES to the TSC conditions (Cinkornpumin *et al*., 2020; Dong *et al*., 2020). Here we have confirmed that transcriptional and epigenetic profiles of naïve-and primed PSCs derived TSCs are highly similar. However, efficient and robust conversion of primed TSCs requires the usage of an induction stage. Application of TSC-m medium that contains GSK3 inhibitor directly on primed cells results in rapid differentiation into non trophoblast somatic lineages **(Figure S5)** thus rendering their derivation very complex and inefficient. Despite this, we have observed, in accordance with (Wei *et al*., 2021), that even direct addition of TSC medium to primed cells does give rise to a small GATA3 positive fraction, from which a normal TSC line can be established **(Figure 5e**). We conclude that application of CHIR, EGF and TGF inhibitor– containing medium on primed, but not naïve hPSCs gives rise both to the trophoblast and to the neural lineage, the latter is capable of more rapid proliferation and quickly overgrow the TSCs.

Wnt/βCatenin signaling is an essential player in the embryonic stem cells field. While mouse ES naïve state demands βCatenin activation, the human naïve conditions mostly require Wnt suppression (Bayerl *et al*., 2021; Guo *et al*., 2017; Theunissen *et al*., 2014; Ying et al., 2008). Moreover, CHIR or Wnt ligands are indispensable components of many hPSCs differentiation protocols. Pax6+Sox1+ population readily emerges in TSC conditions from primed cells but was not visible in converting naïve hPSCs. Our finding might indicate an intriguing difference between naïve and primed hPSCs’ reaction to βCatenin activation that deserves further investigation. Emergence of two very distinct self-renewing populations – TSCs and neural cells - after application of TSC medium on primed cells might also be a subject for further study which should answer, whether primed hPSCs choose their fate in a stochastic manner, or whether there is an intrinsic heterogeneity in the primed hPSC cultures, predisposing some cells to follow trophoblast fate, while the others – the neural one.

YAP translocation to the nuclei of the outer cells of the mouse early blastocyst is an essential trigger for trophoblast lineage specification (Rossant and Tam, 2009). Upregulation of YAP cofactor Tead4 gene in the mouse naïve ES cells converts them to TSCs (Nishioka *et al*., 2009). Moreover, YAP protein shows nuclear localization in the TSCs and prevents cyto-trophoblasts from differentiation (Dong *et al*., 2020; Meinhardt *et al*., 2020). Here we have confirmed that transient overexpression of constitutively active YAP in primed ES cells replaces TGF inhibition in the conversion protocol. Furthermore, YAP knockout hPSCs do not give rise to TSC lines and are degraded at the second passage after induction with TGF inhibitor. Interestingly, some TSC specific markers do show some upregulation upon TGF inhibition in YAP knockout hPSCs, possibly due to the presence of another Hippo signaling effector TAZ. To check whether at least some part of trophoblast program can be induced in complete absence of Hippo signaling effectors, we have attempted to make YAP/TAZ double knockout hPSCs, but failed to do so likely because such double knockout cells do not survive in human primed or naïve PSC supporting conditions. Nevertheless, our results support the conclusion that YAP signaling is sufficient for TSC program induction and necessary for TSCs maintenance.

Conversion of mouse ESCs to TSC is also well established, and implies using transgenes (Nishioka *et al*., 2009; Niwa et al., 2005). This conversion works from naïve ES state, but not from the primed one (Blij *et al*., 2015). This result is believed to reflect higher developmental proximity between naïve ES cells and the trophoblast lineage, since naïve ES cells model preimplantation ICM/epiblast, while the primed cells are more similar to the early post-implantation epiblast (Weinberger *et al*., 2016). The straightforward conversion of human primed ESCs to TSCs suggests that the human primed PSC state still retains a great lineage plasticity to revert into extraembryonic TE fate, thus highlighting a striking difference between human and mouse primed stem cells states.

## Abbreviations used

pdTSC: Primed-derived Trophoblast Stem Cells
ndTSC: Naïve-derived Trophoblast Stem Cells
edTSC: Embryo-derived Trophoblast Stem Cells
hPSC: Human Pluripotent Stem Cells
ESC: Embryonic Stem Cells
hCG: Human Chorionic Gonadotropin
CT: Cytotrophoblasts
STB: Syncytium Trophoblast
EVT: Extravillous Cytotrophoblast
KO: knockout
WT: Wild type

## Acknowledgements

This work was funded by Pascal and Ilana Mantoux; Nella and Leon Benoziyo Center for Neurological Diseases; David and Fela Shapell Family Center for Genetic Disorders Research; Kekst Family Institute for Medical Genetics; Helen and Martin Kimmel Institute for Stem Cell Research; Flight Attendant Medical Research Council (FAMRI); Helen and Martin Kimmel Award for Innovative Investigation; Dr. Beth Rom-Rymer Stem Cell Research Fund; Edmond de Rothschild Foundations; Zantker Charitable Foundation; Estate of Zvia Zeroni; European Research Council (ERC-CoG); Israel Science Foundation (ISF); Minerva; Israel Cancer Research Fund (ICRF) and BSF. We thank the Weizmann Institute management and board for providing critical financial and infrastructural support.

## Author Contribution

S.V. and J.H.H conceived the idea for this project, designed and conducted the experiments and wrote the manuscript. T.S. conducted computational analysis of the data. J.B. assisted with immunohistochemistry experiments, D.S. generated ATAC-seq library. Y.S. assisted with ELF5 methylation experiment. N.N. supervised and conducted computational analysis of the data, and wrote the manuscript. J.H. supervised the project and wrote the paper.

## Figure Legends

**Supplementary Figure 1:**
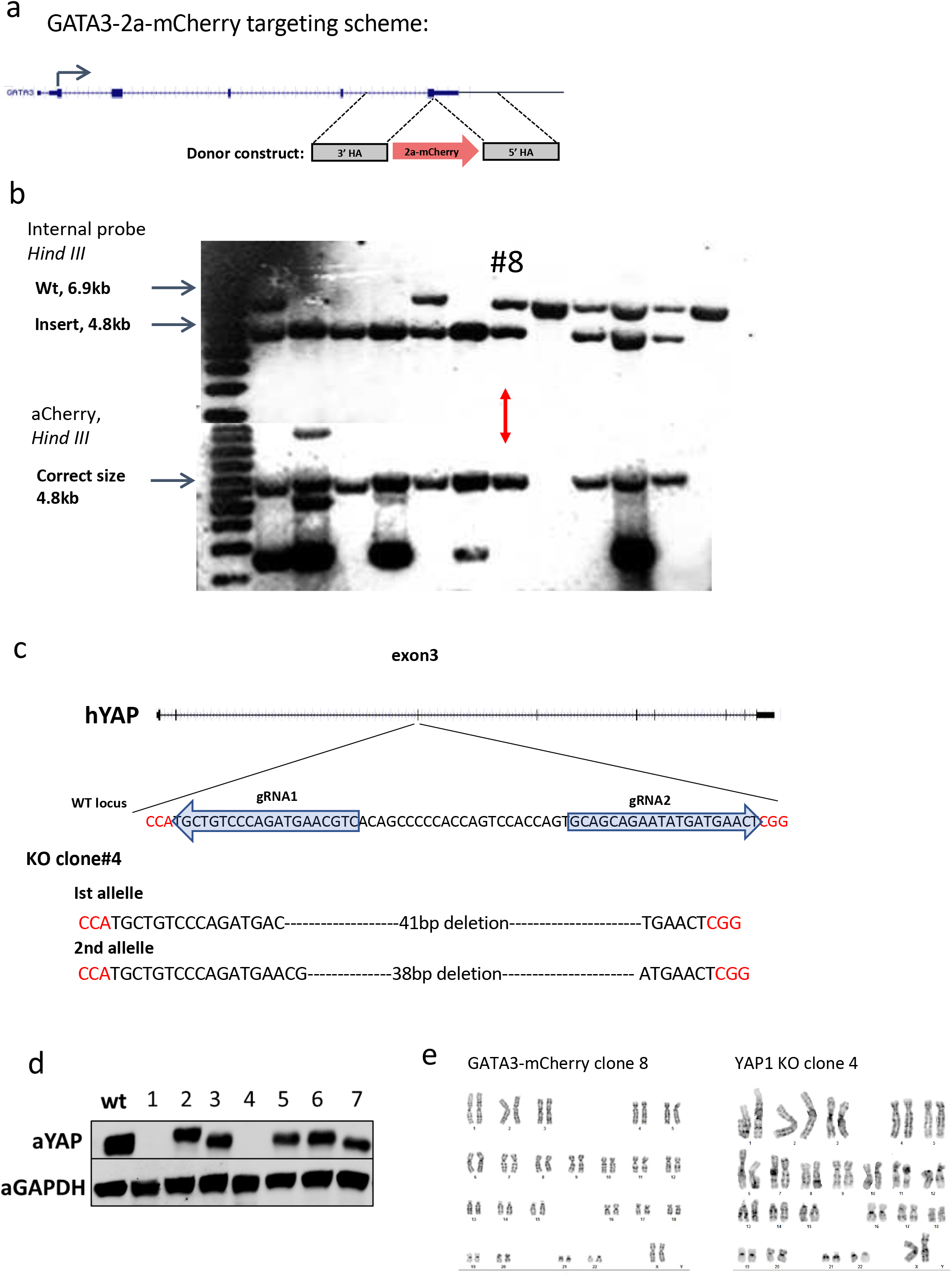
GATA3 and YAP gene targeting. (a) GATA3-2a-mCherry targeting scheme in human PSCs. (b) Southern blot showing for GATA3-2a-mCherry targeted WIBR3 hESC clones. Clone #8 was chosen for further experiments (c) YAP-KO targeting scheme and indels’ sequence in the WIBR3 YAP KO clone. (d) YAP-KO Western blot, showing a successful KO in clones #1 and #4. (e) Normal karyotype of GATA3-mCherry clone #8 and YAP KO clone4.

**Supplementary Figure 2:**
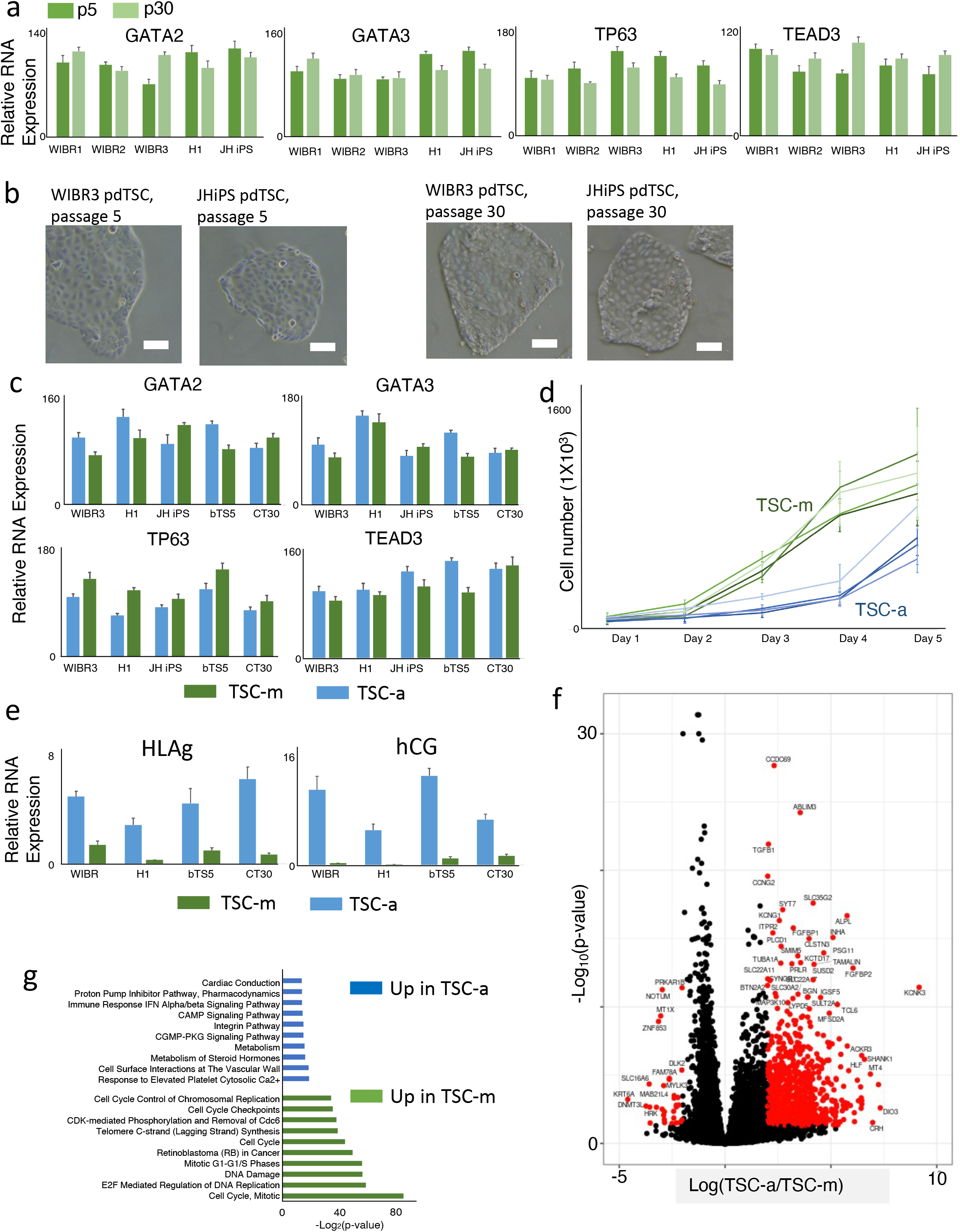
Cellular Identity of pdTSCs. (a) Relative expression of TSC markers GATA2, GATA3, PTGES, DAB2 TP63 & TEAD3 in pdTSCs, 5 and 30 passages after derivation, measured by RT-PCR. (n=3) (b) Bright field images of pdTSC cultures derived from WIBR3 and from JHiPS at passages 5 and 30. Scale bar 100 μm. (c) Relative expression of TSC markers GATA2, GATA3, TP63 & TEAD3 in pdTSCs that were maintained in TSC-a, and TSC-m media for pdTSC lines derived from JHiPS, H1 and WIBR3 as well as bTS5 and CT30 TSCs. (n=3) (d) Growth curve for the cells grown in TSC-a and in TSC-m media, showing a more rapid growth in TSC-m medium. 50×10^3^ cells were seeded at day 0 for each line. (n=3) (e) Upregulation of trophoblast differentiation markers HLA-G and CGb for TSCs grown in TSC-a compared to TSC-m medium. (n=3) (f) Differentially expressed genes between TSC-a and TSC-m maintained pdTSCs. (g) Enriched represented of pathways among differentially expressed genes. Top 10 pathways are presented for each category. Cell cycle genes are induced in TSC-m compared to TSC-a.

**Supplementary Figure 3:**
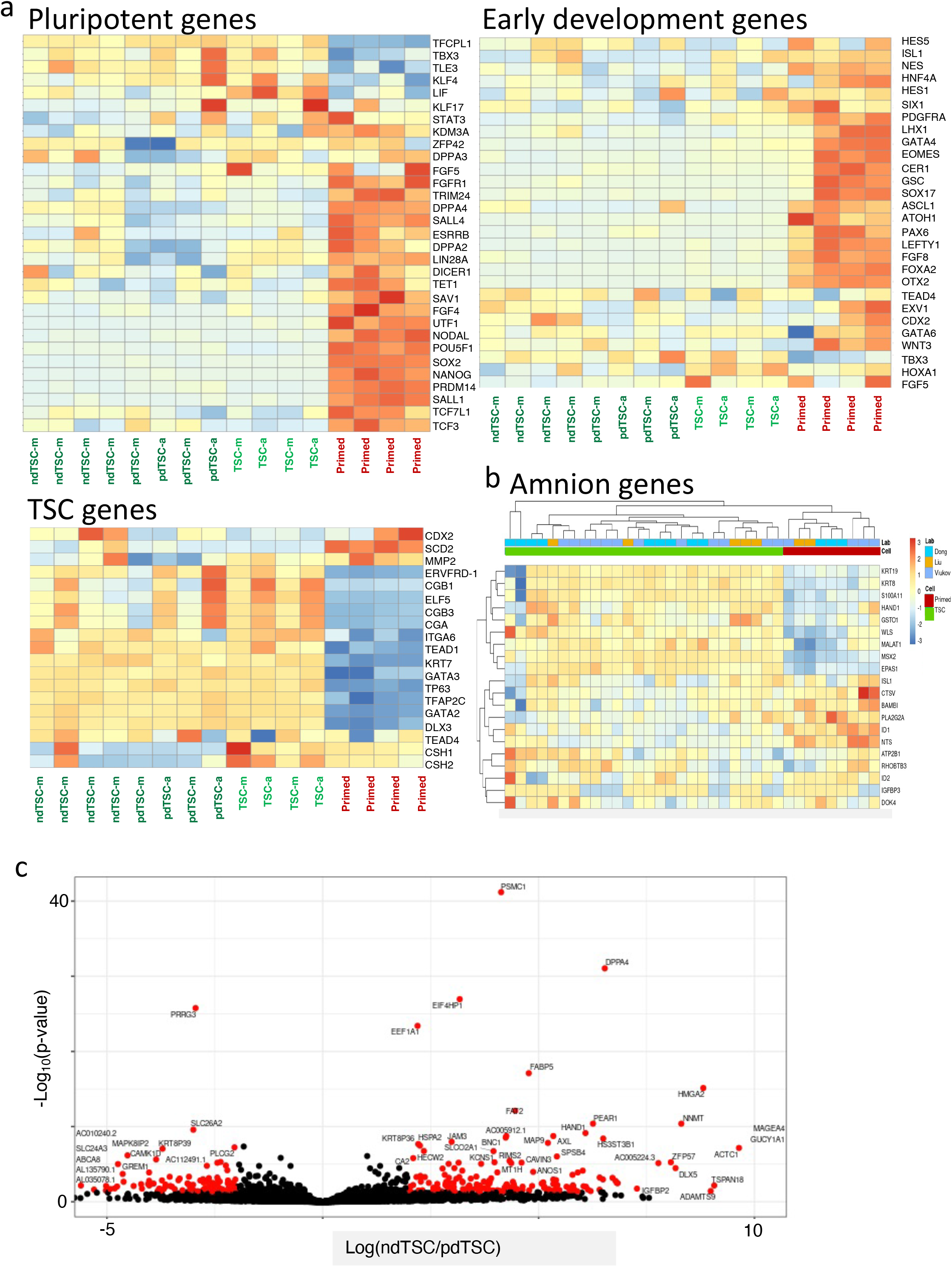
Expression pattern of focused gene groups. (a) Expression pattern of pluripotent genes, early development genes, and TSC genes in our dataset. (b) Expression pattern of amnion genes in our dataset and in external datasets (Dong et al, Liu et al). (c) Differentially expressed genes between TSCs derived from naïve hPSCs and from primed hPSCs.

**Supplementary Figure 4:**
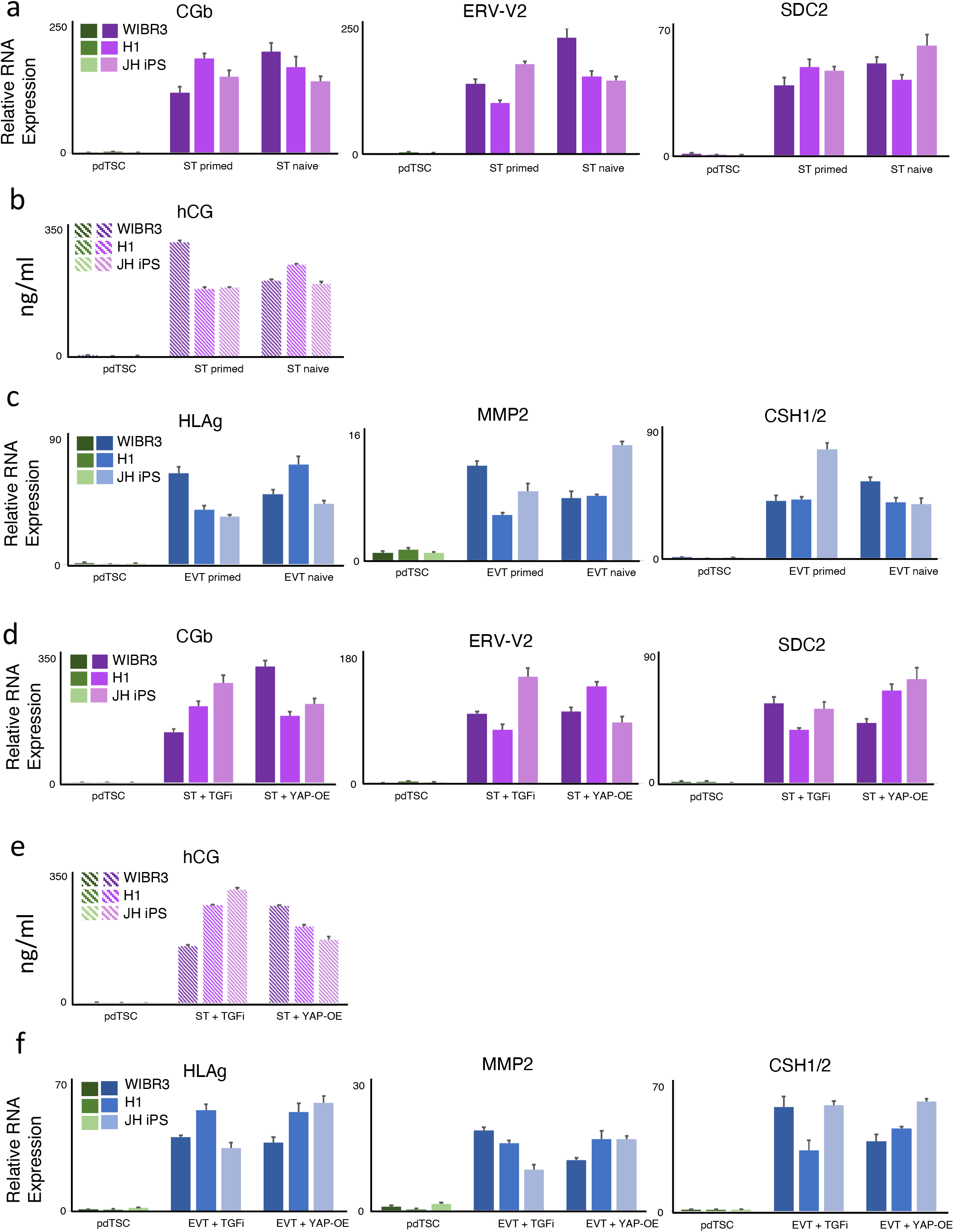
Differentiation assay for TSC derived from Naïve and Primed hPSCs by TGF inhibition and by YAP* overexpression. (a) Relative expression of STB markers CGb, ERV-V2 and SDC2, in TSCs derived from naïve or primed hPSCs and differentiated to STB. (n=3) (b) Secreted CG level for the samples from (a). (n=3) (c) Relative expression of EVT markers HLAG, MMP2, CSH1/2, in TSCs derived from naïve or primed hPSCs and differentiated to EVT. (n=3) (d) Relative expression of ST markers CGb, ERV-V2 and SDC2, in TSCs derived by TGF inhibition or by YAP* overexpression and differentiated to STB. (n=3) (e) Secreted CG level for the samples from (d). (n=3) (f) Relative expression of EVT markers HLAG, MMP2, CSH1/2, in TSCs derived by TGF inhibition or by YAP* overexpression and differentiated to EVT. (n=3)

**Supplementary Figure 5:**
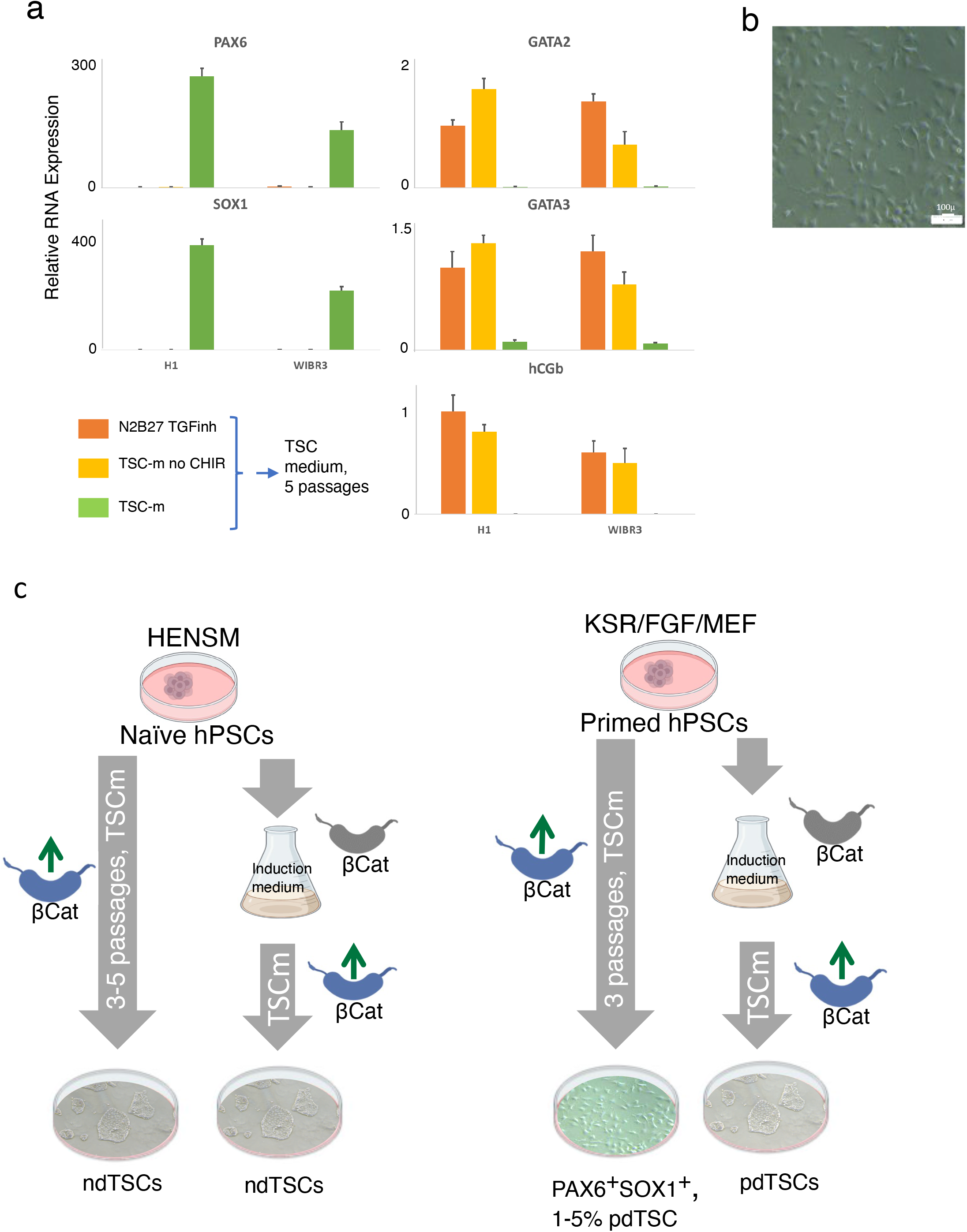
TSC and Neural Cells can be derived from primed PSCs by the same protocol. (a) Expression of neural markers, PAX6 and SOX1, and TSC markers, GATA2, GATA3 and hCGb. H1 and WIBR3 primed hPSC cells were seeded directly in TSC-m medium or stimulated for 5 days with TSC-m medium without CHIR or with TI medium, and then changed to TSC-m. Marker expression was measured at passage 10. (n=5) (b) Bright field image of SOX1^+^ PAX6^+^ positive cells. (c) The proposed scheme: For successful TSC conversion from primed hPSCs bCatenin should not be stimulated for the first days of the derivation protocol, while for naïve hPSCs CHIR addition does not lead to neural identity.

## KEY RESOURCES TABLE

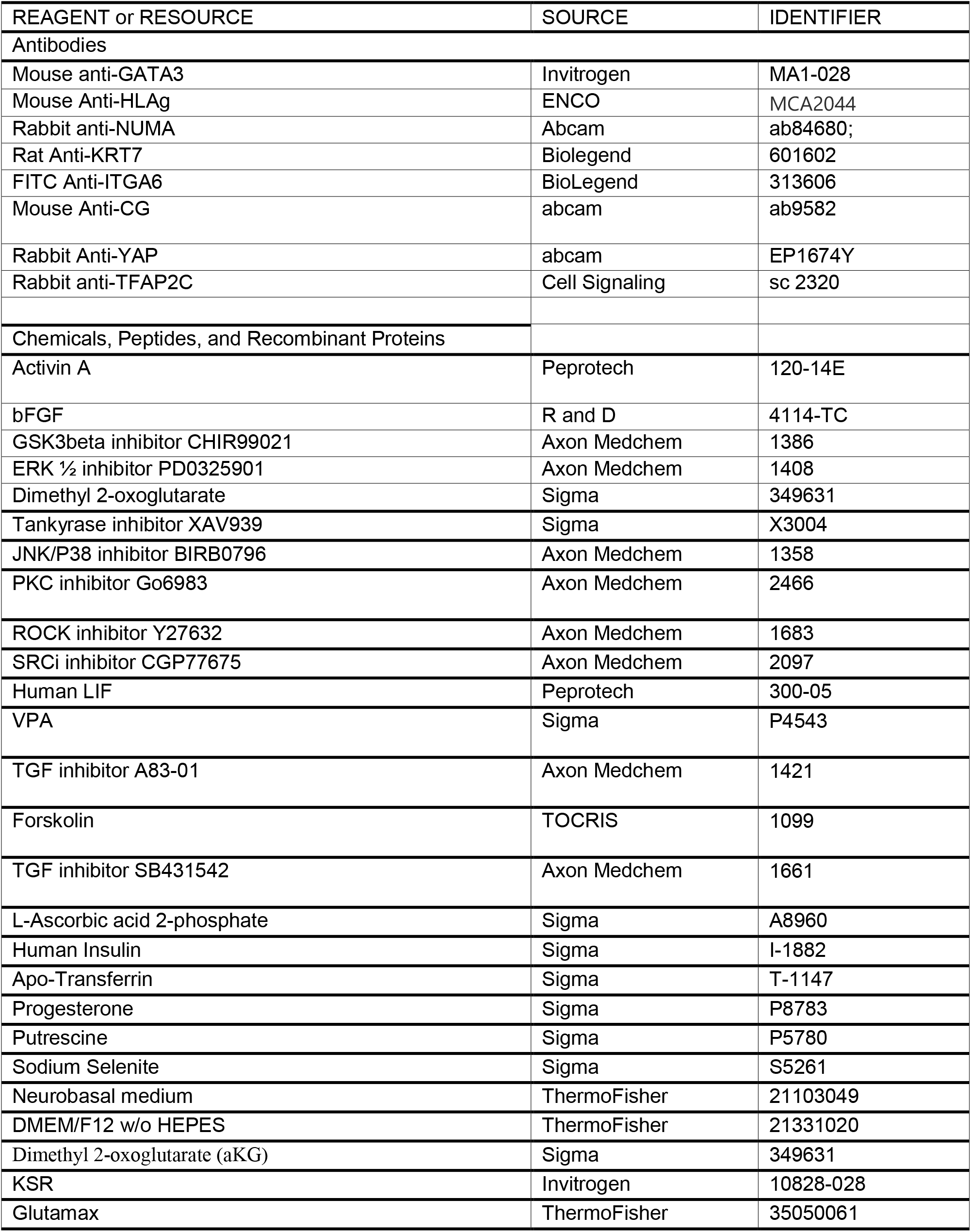
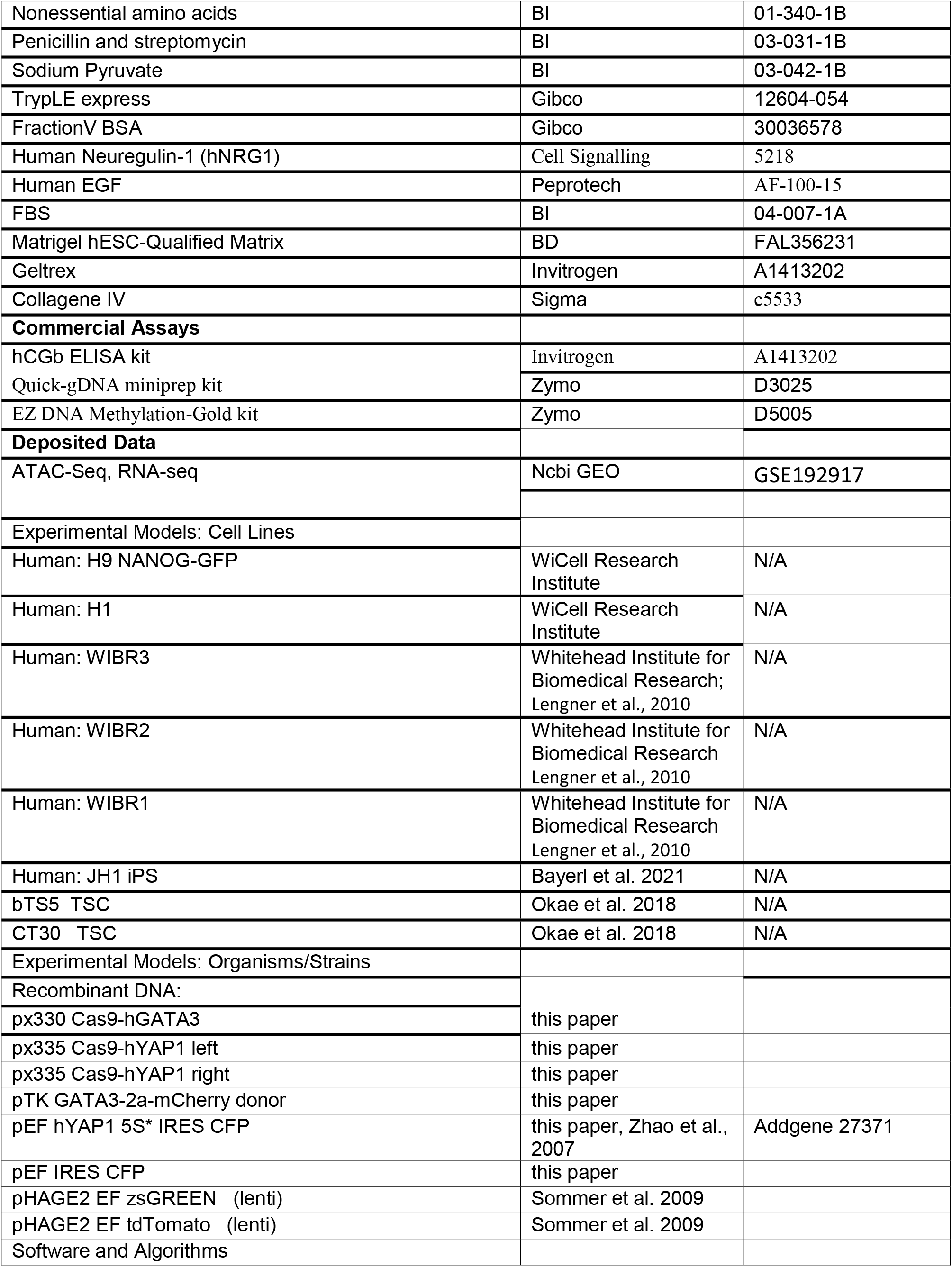
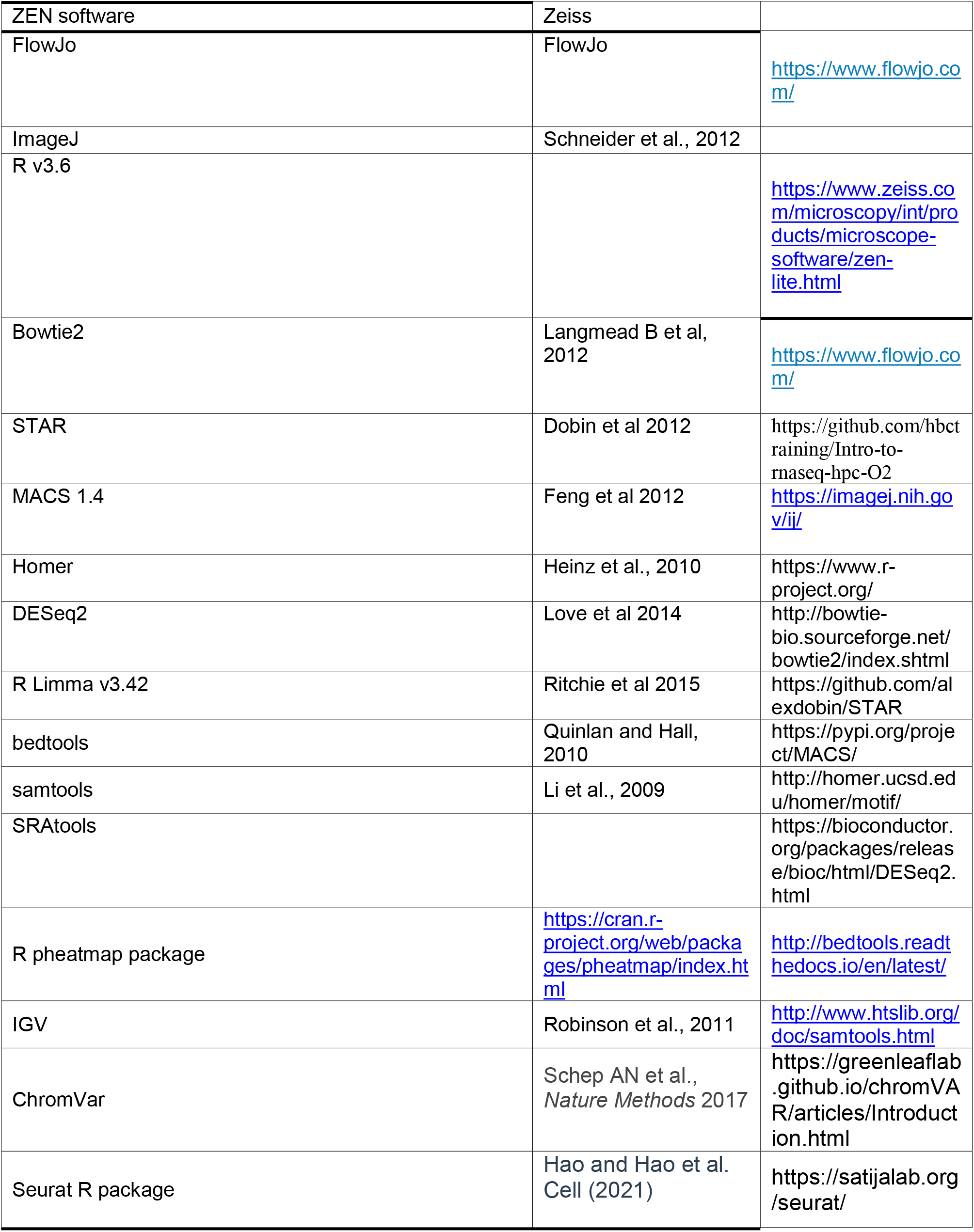

### Lead Contact

Further information and requests for resources and reagents should be directed to the Lead Contact, Jacob Hanna.

### Materials Availability

Unique reagents generated in this study are available from the Lead Contact with a Materials Transfer Agreement.

### Data and Code Availability

All RNA-Seq, ATAC-Seq, data reported in this study have been deposited in Gene Expression Omnibus (https://www.ncbi.nlm.nih.gov/geo/) under Accession code GSE192917. Software/packages used to analyze the dataset are either freely or commercially available.

## Materials and Methods

### Culture of human naïve and primed ESCs and iPSCs

Primed human cells were cultured on irradiated MEFs in DMEM-F12 (Invitrogen 10829) supplemented with 15% Knockout Serum Replacement (Invitrogen 10828-028), 1mM GlutaMAX (Invitrogen), 1% nonessential amino acids (BI 01-340-1B), 1% Penicillin-Streptomycin (BI 03-031-1B), and 8 ng/mL bFGF (Peprotech) in 20% O2 conditions. Naïve cells were cultured in the 5%O2 incubator on the matrigel covered plates, in the HENSM conditions (Bayerl et al., 2021): 240 ml Neurobasal (ThermoFisher 21103049) and 240 ml of DMEM-F12 without HEPES (ThermoFisher 21331020), 5 ml N2 supplement (in-house prepared), 5 mL GlutaMAX (ThermoFisher 35050061), 1% nonessential amino acids (BI 01-340-1B), 1% Penicillin-Streptomycin (BI 03-031-1B), 1% Sodium-pyruvate (BI 03-042-1B), 10 ml B27 supplement (Invitrogen 17504-044), 0.8 mM Dimethyl 2-oxoglutarate (aKG) (Sigma 349631), 1ml Geltrex (Invitrogen A1413202), 50 μg/ml Vitamin C (L-Ascorbic acid 2-phosphate, Sigma A8950), 20 ng/ml recombinant human LIF (Peprotech 300-05), MEKi/ERKi (PD0325901 1 μM - Axon Medchem 1408); WNTi/TNKi (XAV939 2μM - Axon Medchem 1527); P38i/JNKi (BIRB0796 0.8μM - Axon Medchem 1358), PKCi (Go6983 2μM - Axon Medchem 2466), ROCKi Y27632 (1.2μM - Axon Medchem 1683), SRCi (CGP77675 1.2μM – Axon Medchem 2097), hActivin (10ng/ml, Peprotech, 120-14E). All the cells were routinely passaged with 0.05% trypsin. For primed hPSCs 10uM ROCKi was added 24 hrs before and after the passaging, while for naïve PSCs 5uM ROCKi was added for 24 hrs after passaging.

### Targeting GATA3 and YAP1

For making GATA3-mCherry reporter primed WIBR3 hPS cells were electroporated (Bayerl et al. 2021) with the donor construct and px330-GATA3gRNA vector targeting the stop codon. The transfected cells were subcloned and the clones were analyzed by PCR and by Southern Blot with external and anti-mCherry probes. YAP1 gene was knocked out by electroporating WIBR3 WT hES with two Cas9 nickase encoding constructs (px335-YAP1 left and px335-YAP1 right). The clones were analyzed for mutations by PCR and by Western Blot with anti-YAP antibody (Abcam, EP1674Y). To verify the out-of-frame indels the targeted locus was amplified and sequenced.

### pdTSC derivation from primed ES cells and maintenance

Primed human ESCs and iPSCs were trypsinyzed and 10^5^ cells were seeded on matrigel-covered 10cm plates in the induction medium (TI): DMEM/F12 (ThermoFisher 21331020), 1%N2 (in house made), 2% B27, 1% nonessential amino acids (BI 01-340-1B), 1% Penicillin-Streptomycin (BI 03-031-1B), 1% Sodium-pyruvate (BI 03-042-1B), 50 μg/ml Vitamin C (L-Ascorbic acid 2-phosphate, Sigma A8950), 1% GlutaMAX (ThermoFisher 35050061) and 0.5 μM A83-01 (Axon Medchem 1421). The medium was changed daily. After 5 days the cells were passaged on the Collagene IV (Sigma c5533, 5μg/ml) coated plates in the TSC-m medium. TSC-m medium consist of TI medium with addition of 50ng/ml EGF (Peprotech, AF-100-15), 2μM CHIR99021 (Axon Medchem 1386) and 5μM Y27632 (Axon Medchem 1683). The cells were passaged with TrypLE reagent at ratio 1:8 – 1:12 every 3 – 4 days. In some experiments 2μM CHIR99021, 5μM Y27632, 50ng/ml EGF were also added to the TI medium. Maintaining TSC and pdTSC under conventional conditions was performed as described elsewhere (Okae et al., 2018). Briefly, the medium (TSC-a) composed of DMEM/F12 supplemented with 0.1 mM <u>2-mercaptoethanol,</u> 0.2% FBS, 0.5% Penicillin-Streptomycin, 0.3% BSA, 1% ITS-X supplement (homemade), 1.5 μg/ml L-ascorbic acid, 50 ng/ml EGF, 2 μM CHIR99021, 0.5 μM A83-01, 1 μM SB431542, 0.8 mM VPA and 5 μM Y27632. All the cells were grown at 20% O2 with daily medium change. For testing “BAP” conditions we used N2B27 medium supplemented with 25ng/ml BMP4, 0.5μM A83-01 and 1μM PD0325901.

### Growth curve measurement

To compare the growth rate between TSC-m and TSC-a conditions cells were kept in respective media for at least 2 passages and then 50×103 cells were seeded per 1 well of a 6-well plate. The cells were counted daily for 5 days, at that time point all the lines in TSC-m medium have reached confluence. For the growth curve experiment 3 biological replicates were averaged.

### ndTSC derivation from naive ES cells

For derivation of ndTSC lines from naïve conditions the cells growing in HENSM medium were trypsinized and seeded on matrigel-covered 10cm plates at the density of 1×10^5^ cells per plate either into TI medium or into TSC-m medium. Around day 5 the cells were passaged at the ratio of 1:2 – 1:4 to Collagene IV coated plates into TSC-m medium. Subsequently the cells were passaged every 3-4 days, and stable cultures with homogeneous TSC morphology and >90% cells expressing GATA3 protein were established by passage 3-5 depending on the background.

### ESC and induced TSC karyotyping

Chromosome analysis was performed by G-banding following standard procedure. Briefly, 50% confluent cells were treated with 100ng/ml colcemid for 40 min, Trypsinized, treated with hypotonic solution, fixed and applied on slides. The slides were stained with Giemsa stain (Sigma GS500). The metaphase spreads were analyzed according to the International System for Human Cytogenetic Nomenclature. At least 20 cells were fully analyzed for each cell line.

### ST and EVT differentiation assay

TSC differentiation and engraftment were performed exactly as described (Okae et al., 2018). Breifly, for ST differentiation the cells were seeded in the 6-well collagen IV precoated plates at the density of 10^5^ cells per well in the ST(2D) medium (DMEM/F12, 0.1mM 2-mercaptoethanol, 0.5% penicillin-streptomycine, 1% ITS-X supplement, 0.3% BSA, 4% KSR, 2.5uM Y27632 and 2uM forskolin). The medium was changed at day 2 and 4 and the cells were analyzed 6 days after plating. For measuring secreted hCG level the supernatant was collected and the hCG concentration was measured with hCG ELISA kit. For control the supernatants from 70% confluent TS cells, left without medium change for 2 days, were used. For EVT differentiation 105 TS cells were seeded per 1 well of a Col IV precoated 6-well plate in the EVT medium (DMEM/F12, 0.1μM 2-mercaptoethanol, 0.5% penicillin-streptomycine, 1% ITS-X supplement, 0.3% BSA, 4% KSR, 100ng/ml NRG1, 2.5μM Y27632, 7.5μM A83-01 supplemented with 2% Matrigel). 3 days after plating the medium was switched to EVT with 0.5% Matrigel, without NRG1. 6 days after plating the cells were passaged at 1:2 – 1:3 ratio to the EVT medium without NRG1 and KSR with 0.5% Matrigel. 2 days later the cells were analyzed.

### Lesion formation in the NOD-SCID mice

For engraftment of the cells into NOD-SCID mice 10^7^ induced TSCs grown in TSC-m medium were dissociated, resuspended in 200uL of 2:1 mixture of DMEM/F12-N2B27 and Matrigel, and injected subcutaneously into 6 weeks old male mice. Lesions were dissected and serum collected 10 days after injection. hCG level in the serum was measured using hCG ELISA kit.

### RT-PCR analysis

Total RNA was isolated using Trizol reagent (Ambion life technologies). 1μg of total RNA was reverse transcribed using a High-Capacity Reverse Transcription Kit (Applied Biosystems). Quantitative PCR analysis was performed in triplicate using 1/100 of the reverse transcription reaction in a Viia7 platform (Applied Biosystems) with the relevant primers (Table 1). Normalization to RPL3 and HPRT was done across all experiments, and data is presented in relevance to the indicated sample set as 1.

### Statistics

Error bars indicate standard deviation of independent experiments, as indicated by “n” in the figure legend.

### Immunofluorescence staining

Cells growing on glass cover slips (13mm 1.5H; Marienfeld, 0117530) were fixed with 4% paraformaldehyde/phosphate buffer for 10 min at room temperature, washed three times with PBS, and permeabilized in PBS/0.1% Triton for 10 min. Cells were blocked with blocking solution (2% normal goat serum, 0.1% BSA in PBS/0.05% Tween) for 1h at RT and incubated with primary antibody diluted in blocking solution overnight at 4°. Cells were then washed three times with PBS/0.05% Tween, incubated with secondary antibodies for 1 hour at room temperature, washed in PBS/0.05% Tween, counterstained with DAPI (1 μg/ml), washed again three times with PBS/Tween 0.05%, mounted with Shandon Immu-Mount (Thermo Scientific, 9990412), and imaged. All comparative experiments were done simultaneously.

The lesions from the NOD-SCID mice were fixated overnight in 4% PFA, washed three times in PBS and dehydrated in sequential Ethanol dilutions. After standard paraffinization and embedding, tissue blocks were sectioned on a microtome and mounted onto Superfrost plus slides (Thermo Scientific, Menzel-Glaser), dried at 38º C for 4 hours and stored at 4º C until further processing. The sections were stained with H&E. For staining with antibodies the sections were rehydrated, antigen retrieved in Sodium citrate buffer and pressure cooker, rinsed in PBS and treated with 0.3% H2O2 to reduce background staining. After permeabilization in 0.1% Triton X-100 in PBS three times for 2 min., samples were blocked in 10% normal donkey serum in PBS in humidified chamber for 20 min. at RT. Slides were then incubated in the appropriate primary antibody diluted in 1% BSA in 0.1% Triton X-100 at 4 °C overnight. Sections were then washed three times (5 min each) in washing buffer (0.1% Triton X-100 in PBS) incubated with appropriate fluorochrome-conjugated secondary antibodies diluted in 1% BSA in 0.1% Triton X-100 at RT for 1 h in the dark. For human cell specific NUMA staining, signal enhancement was performed by usage of Biotin-SP conjugated 2nd antibody for 30 min and subsequent incubation with Streptavidin-Cy3 antibody for 30 min. All sections were washed three times in washing buffer for 10 min each, counter stained with DAPI for 20 min, rinsed twice in washing buffer and mounted with Shandon Immuno-Mount (Thermo Scientific, 9990412).

The following antibodies were use at the indicated dilutions: Mouse anti-GATA3 (Invitrogen, MA1-028, 1:200), Mouse Anti-HLAg (ENCO, MCA2044, 1:200), Rabbit Anti-NUMA (Abcam, ab84680, 1:100), Rat Anti-Krt7 (1:200), Mouse Anti-CG (abcam ab84680 1:200), Rabbit anti-YAP,(abcam EP1674Y, 1:200), Rabbit anti TFAP2C (Cell Signalling, 2320).

### Methylation analysis of ELF5 promoter

DNA was isolated from cells using the Quick-gDNA miniprep kit (Zymo). DNA (50ng) was then converted by bisulfite using the EZ DNA Methylation-Gold kit (Zymo). [-136 - +238] region of ELF5 promoter was amplified by sequential PCR using nested primers (Table1) and the PCR product was cloned into pJET2.1 vector. 10 bacterial clones were Sanger sequenced for each sample.

### ATACseq

Cells were trypsinized and counted, 50,000 cells were centrifuged at 500*g* for 3 min, followed by a wash using 50 μl of cold PBS and centrifugation at 500*g* for 3 min. Cells were lysed using cold lysis buffer (10 mM Tris-HCl, pH 7.4, 10 mM NaCl, 3 mM MgCl_2_ and 0.1% IGEPAL CA-630). Immediately after lysis, nuclei were spun at 500*g* for 10 min using a refrigerated centrifuge. Next, the pellet was resuspended in the transposase reaction mix (25 μl 2× TD buffer, 2.5 μl transposase (Illumina) and 22.5 μl nuclease-free water). The transposition reaction was carried out for 30 min at 37 °C and immediately put on ice. Directly afterwards, the sample was purified using a Qiagen MinElute kit. Following purification, the library fragments were amplified using custom Nextera PCR primers 1 and 2 for a total of 12 cycles. Following PCR amplification, the libraries were purified using a QiagenMinElute Kit and sequenced.

### RNA-seq library preparation

Total RNA was extracted from indicated cell lines at 50% confluency with Trizol reagent. RNA was next utilized for RNA-Seq by ScriptSeq Preparation Kit v2 (Illumina) according to manufacturer’s instruction.

### RNA-seq analysis

Overall, 18 samples were sequenced using poly-A single-end and paired-end strategy (4 Primed hPSCs, 2 fibroblasts, 4 TSC and 4 pdTSCs and 4 ndTSCs, **Supplementary Table 2**). STAR software version 2.5.3a was used to align reads to human GRCh38 reference genome (2013), using the following flags: --outFilterMultimapNmax 1 --outReadsUnmapped Fastx --twopassMode Basic --outSAMstrandField intronMotif. Read count values were estimated with HTSeq V0.7.2 software over all aligned reads using GRCh38 general feature format (GFF), with the following flags: -a 10 -s no -t exon -i gene_id. Mitochondrial and ribosomal genes were filtered out. Genes with a at least 2 counts > 5 were kept for further analysis. Normalization of counts and detection of differentially expressed genes were done using R DESeq2 V1.26 software. Differentially expressed genes between TSC and Primed samples were selected by log2(Fold change)> 2 or log2(Fold change) < -2, and FDR-adjusted p-value< 0.05. Functional enrichment analysis was done with GeneAnalytics software (https://www.genecards.org/). Clustering with previously published data was done as follows: raw data were downloaded from GSE150616 (Liu *et al*., 2020) and GSE138762 (Dong *et al*., 2020), and undergone the same alignment and filtering as described above. Unified data were corrected for batch-effects using limma R package. PCA analysis and plotting was performed by R V3.6. Data is available as ncbi GEO series number GSE192917.

### ATAC-seq library

Cells were trypsinized and counted, 50,000 cells were centrifuged at 500*g* for 3 min, followed by a wash using 50 μl of cold PBS and centrifugation at 500*g* for 3 min. Cells were lysed using cold lysis buffer (10 mM Tris-HCl, pH 7.4, 10 mM NaCl, 3 mM MgCl_2_ and 0.1% IGEPAL CA-630). Immediately after lysis, nuclei were spun at 500*g* for 10 min using a refrigerated centrifuge. Next, the pellet was resuspended in the transposase reaction mix (25 μl 2× TD buffer, 2.5 μl transposase (Illumina) and 22.5 μl nuclease-free water). The transposition reaction was carried out for 30 min at 37 °C and immediately put on ice. Directly afterwards, the sample was purified using a Qiagen MinElute kit. Following purification, the library fragments were amplified using custom Nextera PCR primers 1 and 2 for a total of 12 cycles. Following PCR amplification, the libraries were purified using a QiagenMinElute Kit and sequenced.

### ATAC-seq analysis

Overall, 17 new samples were sequenced using ATAC-seq strategy (3 Primed hESCs, 4 TSC and 4 pdTSCs and 6 ndTSC, **Supplementary table 2**).

Reads were aligned to hg38 human genome using Bowtie2 with the parameter -X2000 (allowing fragments up to 2 kb to align). R ChromVar package was used to identify motifs that are differentially represented, and to cluster the samples according to their motif enrichment pattern.

### Projection onto single-cell data

The projection of our RNA-seq samples on single-cell data was done as following: single-cell data was downloaded from previous publications (Tyser et al., 2020; Zheng et al., 2019). Counts were transferred to R Seurat object, normalized anchored and integrated into a single UMAP. Variable features from this integration were used to filter our batch RNA-seq samples. The single-cell combined dataset and the batch RNA-seq data were normalized separately, and then integrated and rescaled using linear regression.

## References

Amita, M., Adachi, K., Alexenko, A.P., Sinha, S., Schust, D.J., Schulz, L.C., Roberts, R.M., and Ezashi, T. (2013). Complete and unidirectional conversion of human embryonic stem cells to trophoblast by BMP4. Proc Natl Acad Sci U S A 110, E1212–1221. 10.1/073pnas.1303094110.

Avraham, R., and Yarden, Y. (2011). Feedback regulation of EGFR signalling: decision making by early and delayed loops. Nat Rev Mol Cell Biol 12, 104–117. 10.1038/nrm3048.

Bayerl, J., Ayyash, M., Shani, T., Manor, Y.S., Gafni, O., Massarwa, R., Kalma, Y., Aguilera-Castrejon, A., Zerbib, M., Amir, H., et al. (2021). Principles of signaling pathway modulation for enhancing human naive pluripotency induction. Cell Stem Cell. 10.1016/j.stem.2021.04.001.

Bhattacharya, S., Mukherjee, A., Pisano, S., Altshuler, A., Nasser, W., Dey, S., Kaganovsky, A., Amitai-Lange, A., Mimouni, M., Socea, S., et al. (2021). Soft limbal niche maintains stem cell compartmentalization and function through YAP. bioRxiv, 2021.2005.2025.445490. 10.1101/2021.05.25.445490.

Blij, S., Parenti, A., Tabatabai-Yazdi, N., and Ralston, A. (2015). Cdx2 efficiently induces trophoblast stem-like cells in naive, but not primed, pluripotent stem cells. Stem Cells Dev 24, 1352–1365. 10.1089/scd.2014.0395.

Chaikuad, A., and Bullock, A.N. (2016). Structural Basis of Intracellular TGF-β Signaling: Receptors and Smads. Cold Spring Harb Perspect Biol 8. 10.1101/cshperspect.a022111.

Cinkornpumin, J.K., Kwon, S.Y., Guo, Y., Hossain, I., Sirois, J., Russett, C.S., Tseng, H.W., Okae, H,.Arima, T., Duchaine, T.F., et al. (2020). Naive Human Embryonic Stem Cells Can Give Rise to Cells with a Trophoblast-like Transcriptome and Methylome. Stem Cell Reports 15, 198–213. 10.1016/j.stemcr.2020.06.003.

Cornacchia, D., Zhang, C., Zimmer, B., Chung, S.Y., Fan, Y., Soliman, M.A., Tchieu, J., Chambers, S.M., Shah, H., Paull, D., et al. (2019). Lipid Deprivation Induces a Stable, Naive-to-Primed Intermediate State of Pluripotency in Human PSCs. Cell Stem Cell 25, 120–136 e110. 10.1016/j.stem.2019.05.001.

Dong, C., Beltcheva, M., Gontarz, P., Zhang, B., Popli, P., Fischer, L.A., Khan, S.A., Park, K.M., Yoon, E.J., Xing, X., et al. (2020). Derivation of trophoblast stem cells from naive human pluripotent stem cells. Elife 9. 10.7554/eLife.52504.

Guo, G., Stirparo, G.G., Strawbridge, S.E., Spindlow, D., Yang, J., Clarke, J., Dattani, A., Yanagida, A., Li, M.A., Myers, S., et al. (2021). Human naive epiblast cells possess unrestricted lineage potential. Cell Stem Cell 28, 1040–1056 e1046. 10.1016/j.stem.2021.02.025.

Guo, G., von Meyenn, F., Rostovskaya, M., Clarke, J., Dietmann, S., Baker, D., Sahakyan, A., Myers, S., Bertone, P., Reik, W., et al. (2017). Epigenetic resetting of human pluripotency. Development 144, 2748–2763. 10.1242/dev.146811.

Hemberger, M., Udayashankar, R., Tesar, P., Moore, H., and Burton, G.J. (2010). ELF5-enforced transcriptional networks define an epigenetically regulated trophoblast stem cell compartment in the human placenta. Hum Mol Genet 19, 2456–2467. 10.1093/hmg/ddq128.

Io, S., Kabata, M., Iemura, Y., Semi, K., Morone, N., Minagawa, A., Wang, B., Okamoto, I., Nakamura, T., Kojima, Y., et al. (2021). Capturing human trophoblast development with naive pluripotent stem cells in vitro. Cell Stem Cell 28, 1023–1039 e1013. 10.10/16j.stem.2021.03.013.

James, J.L., Carter, A.M., and Chamley, L.W. (2012). Human placentation from nidation to 5 weeks of gestation. Part I: What do we know about formative placental development following implantation? Placenta 33, 327–334. 10.1016/j.placenta.2012.01.020.

Latos, P.A., and Hemberger, M. (2016). From the stem of the placental tree: trophoblast stem cells and their progeny. Development 143, 3650–3660. 10.1242/dev.133462.

Lengner, C.J., Gimelbrant, A.A., Erwin, J.A., Cheng, A.W., Guenther, M.G., Welstead, G.G., Alagappan, R., Frampton, G.M., Xu, P., Muffat, J., et al. (2010). Derivation of pre-X inactivation human embryonic stem cells under physiological oxygen concentrations. Cell 141, 872–883. 10.1016/j.cell.2010.04.010.

Liu, X., Ouyang, J.F., Rossello, F.J., Tan, J.P., Davidson, K.C., Valdes, D.S., Schroder, J., Sun, Y.B.Y., Chen, J., Knaupp, A.S., et al. (2020). Reprogramming roadmap reveals route to human induced trophoblast stem cells. Nature 586, 101–107. 10.1038/s41586-020-2734-6.

Liu, X., Tan, J.P., Schroder, J., Aberkane, A., Ouyang, J.F., Mohenska, M., Lim, S.M., Sun, Y.B.Y., Chen, J., Sun, G., et al. (2021). Modelling human blastocysts by reprogramming fibroblasts into iBlastoids. Nature 591, 627–632. 10.1038/s41586-021-03372-y.

Meinhardt, G., Haider, S., Kunihs, V., Saleh, L., Pollheimer, J., Fiala, C., Hetey, S., Feher, Z., Szilagyi, A., Than, N.G., and Knofler, M. (2020). Pivotal role of the transcriptional co-activator YAP in trophoblast stemness of the developing human placenta. Proc Natl Acad Sci U S A 117, 13562–13570. 10.1073/pnas.2002630117.

Mischler, A., Karakis, V., Mahinthakumar, J., Carberry, C.K., San Miguel, A., Rager, J.E., Fry, R.C., and Rao, B.M. (2021). Two distinct trophectoderm lineage stem cells from human pluripotent stem cells. J Biol Chem 296, 100386. 10.1016/j.jbc.2021.100386.

Niakan, K.K., Han, J., Pedersen, R.A., Simon, C., and Pera, R.A. (2012). Human pre-implantation embryo development. Development 139, 829–841. 10.1242/dev.060426.

Nishioka, N., Inoue, K., Adachi, K., Kiyonari, H., Ota, M., Ralston, A., Yabuta, N., Hirahara, S., Stephenson, R.O., Ogonuki, N., et al. (2009). The Hippo signaling pathway components Lats and Yap pattern Tead4 activity to distinguish mouse trophectoderm from inner cell mass. Dev Cell 16, 398–410. 10.1016/j.devcel.2009.02.003.

Niwa, H., Toyooka, Y., Shimosato, D., Strumpf, D., Takahashi, K., Yagi, R., and Rossant, J. (2005). Interaction between Oct3/4 and Cdx2 determines trophectoderm differentiation. Cell 123, 917-929. 10/1016.j.cell.2005.08.040.

Okae, H., Toh, H., Sato, T., Hiura, H., Takahashi, S., Shirane, K., Kabayama, Y., Suyama, M., Sasaki, H., and Arima, T. (2018). Derivation of Human Trophoblast Stem Cells. Cell Stem Cell 22, 50–63 e56. 10.1016/j.stem.2017.11.004.

Pefani, D.E., Pankova, D., Abraham, A.G., Grawenda, A.M., Vlahov, N., Scrace, S., and E, O.N. (2016). TGF-β Targets the Hippo Pathway Scaffold RASSF1A to Facilitate YAP/SMAD2 Nuclear Translocation. Mol Cell 63, 156–166. 10.1016/j.molcel.2016.05.012.

Petropoulos, S., Edsgard, D., Reinius, B., Deng, Q., Panula, S.P., Codeluppi, S., Plaza Reyes, A., Linnarsson, S., Sandberg, R., and Lanner, F. (2016). Single-Cell RNA-Seq Reveals Lineage and X Chromosome Dynamics in Human Preimplantation Embryos. Cell 165, 1/10.1016.012-1026j.cell.2016.03.023.

Rossant, J., and Tam, P.P. (2009). Blastocyst lineage formation, early embryonic asymmetries and axis patterning in the mouse. Development 136, 701–713. 10.1242/dev.017178.

Soncin, F., Khater, M., To, C., Pizzo, D., Farah, O., Wakeland, A., Arul Nambi Rajan, K., Nelson, K.K., Chang, C.W., Moretto-Zita, M., et al. (2018). Comparative analysis of mouse and human placentae across gestation reveals species-specific regulators of placental development. Development 145. 10/1242.dev.156273.

Theunissen, T.W., Powell, B.E., Wang, H., Mitalipova, M., Faddah, D.A., Reddy, J., Fan, Z.P., Maetzel, D., Ganz, K., Shi, L., et al. (2014). Systematic identification of culture conditions for induction and maintenance of naive human pluripotency. Cell Stem Cell 15, 471–487. 10.1016/j.stem.2014.07.002.

Turco, M.Y., Gardner, L., Kay, R.G., Hamilton, R.S., Prater, M., Hollinshead, M.S., McWhinnie, A., Esposito, L., Fernando, R., Skelton, H., et al. (2018). Trophoblast organoids as a model for maternal-fetal interactions during human placentation. Nature 564, 263–267. 10.1038/s41586-018-0753-3.

Turco, M.Y., and Moffett, A. (2019). Development of the human placenta. Development 146. 10.1242/dev.163428.

Tyser, R.C.V., Mahammadov, E., Nakanoh, S., Vallier, L., Scialdone, A., and Srinivas, S. (2020). A spatially resolved single cell atlas of human gastrulation. bioRxiv, 2020.2007.2021.213512. 10.1101/2020.07.21.213512.

Varelas, X., Miller, B.W., Sopko, R., Song, S., Gregorieff, A., Fellouse, F.A,. Sakuma, R., Pawson, T., Hunziker, W., McNeill, H., et al. (2010). The Hippo pathway regulates Wnt/beta-catenin signaling. Dev Cell 18, 579–591. 10.1016/j.devcel.2010.03.007.

Wei, Y., Wang, T., Ma, L., Zhang, Y., Zhao, Y., Lye, K., Xiao, L., Chen, C., Wang, Z., Ma, Y., et al. (2021). Efficient derivation of human trophoblast stem cells from primed pluripotent stem cells. Sci Adv 7. 10.1126/sciadv.abf4416.

Weinberger, L., Ayyash, M., Novershtern, N., and Hanna, J.H. (2016). Dynamic stem cell states: naive to primed pluripotency in rodents and humans. Nat Rev Mol Cell Biol 17, 155–169. 10.1038/nrm.2015.28.

Ying, Q.L., Wray, J., Nichols, J., Batlle-Morera, L., Doble, B., Woodgett, J., Cohen, P., and Smith, A. (2008). The ground state of embryonic stem cell self-renewal. Nature 453, 519–523. 10.1038/nature06968.

Yu, L., Wei, Y., Duan, J., Schmitz, D.A., Sakurai, M., Wang, L., Wang, K., Zhao, S., Hon, G.C., and Wu, J. (2021). Blastocyst-like structures generated from human pluripotent stem cells. Nature 591. 620-626, 10.1038s41586-021-03356-y.

Zhao, B., Wei, X., Li, W., Udan, R.S., Yang, Q., Kim, J., Xie, J., Ikenoue, T., Yu, J., Li, L., et al. (2007). Inactivation of YAP oncoprotein by the Hippo pathway is involved in cell contact inhibition and tissue growth control. Genes Dev 21, 2747–2761. 10.1101/gad.1602907.

Zhao, C., Reyes, A.P., Schell, J.P., Weltner, J., Ortega, N., Zheng, Y., Björklund, Å.K., Rossant, J., Fu, J., Petropoulos, S., and Lanner, F. (2021). Reprogrammed iBlastoids contain amnion-like cells but not trophectoderm. bioRxiv, 2021.2005.2007.442980. 10.1101/2021.05.07.442980.

Zheng, Y., Xue, X., Shao, Y., Wang, S., Esfahani, S.N., Li, Z., Muncie, J.M., Lakins, J.N., Weaver, V.M., Gumucio, D.L., and Fu, J. (2019). Controlled modelling of human epiblast and amnion development using stem cells. Nature 573, 421–425. 10.1038/s41586-019-1535-2.

